# Enhancing DNA metabarcoding performance and applicability with bait capture enrichment and DNA from conservative ethanol

**DOI:** 10.1101/580464

**Authors:** M. Gauthier, L. Konecny-Dupré, A. Nguyen, V. Elbrecht, T. Datry, C.J. Douady, T. Lefébure

## Abstract

Metabarcoding is often presented as an alternative identification tool to compensate for coarse taxonomic resolution and misidentification encountered with traditional morphological approaches. However, metabarcoding comes with two major impediments which slow down its adoption. First, the picking and destruction of organisms for DNA extraction are time and cost consuming and do not allow organism conservation for further evaluations. Second, current metabarcoding protocols include a PCR enrichment step which induces errors in the estimation of species diversity and relative biomasses. In this study, we first evaluated the capacity of capture enrichment to replace PCR enrichment using controlled freshwater macrozoobenthos mock communities. Then, we tested if DNA extracted from the fixative ethanol (etDNA) of the same mock communities can be used as an alternative to DNA extracted from pools of whole organisms (bulk DNA). We show that capture enrichment provides more reliable and accurate representation of species occurrences and relative biomasses in comparison with PCR enrichment for bulk DNA. While etDNA does not permit to estimate relative biomasses, etDNA and bulk DNA provide equivalent species detection rates. Thanks to its robustness to mismatches, capture enrichment is already an efficient alternative to PCR enrichment for metabarcoding and, if coupled to etDNA, is a time-saver option in studies where presence information only is sufficient.

## INTRODUCTION

Reliable and accurate taxa identification is fundamental in biological sciences. Poor taxonomic identification can lead to cascades of error affecting our knowledge and understanding not only in theoretical and fundamental biology, but also in applied fields leading to poor management decisions (Bortolus, 2008). It distorts our ability to infer processes in ecology and evolution, to manage and conserve human-impacted systems and to carry out human health and resource programs (Bortolus, 2008; Leys, Keller, Räsänen, Gattolliat, & Robinson, 2016; Prié, Puillandre, & Bouchet, 2013). Poor taxonomic identification occurs when identification to species level is not possible (coarse taxonomic resolution) or when identification is incorrect (misidentification). For example, Martin, Adamowicz, and Cottenie (2016) investigated macrozoobenthos community distribution in freshwater streams at different taxonomic level (family, genus and species) and found a spatial structure only when the species identification level was reached. In Vietnam, misidentification in *Anopheles* species lead to mismanagement in control program of the Malaria disease as the main target species was a non-vector one (Van Bortel et al., 2001). This kind of difficulties leading to poor taxonomic identification and thus to misinterpretation and erroneous conclusions is often associated to morphological based taxonomy (Baird & Hajibabaei, 2012; Creer et al., 2016). Cryptic species, limited expertise, damaged or juvenile specimens and even cost and time constraints in the case of applied research are all problems encountered in morphological identification leading to misidentification or coarse taxonomic resolution (Bringloe, Cottenie, Martin, & Adamowicz, 2016; Hajibabaei, Baird, Fahner, Beiko, & Golding, 2016; Ji et al., 2013; Sweeney et al., 2011).

In the past decade, DNA-based identification has been proposed as an alternative to morphological approaches. DNA barcoding uses the sequence of a genetic marker of one specimen, hereafter called a DNA barcode, usually of an organelle genome for eukaryotes (e.g. mitochondria for animals and chloroplast for plants, see Creer et al. (2016) for overview) and assign it to a species name within a reference database (Hebert, Ratnasingham, & de Waard, 2006). DNA barcoding supposedly addresses limitations in morphological identification by accurately discriminating species regardless of their morphology (Sweeney et al., 2011), development stages (Hubert, Delrieu-Trottin, Irisson, Meyer, & Planes, 2010) or sex (Forshaw, 2010). Recent advances in high throughput technologies enabled the emergence of DNA metabarcoding, the barcoding of pool of specimens in a single reaction which permit to work on whole community at once (Pompanon et al, 2011). Metabarcoding has already been adopted as a routine identification tool in microbial community ecology (Abdelfattah, Malacrinò, Wisniewski, Cacciola, & Schena, 2018). In macro-organism community studies, the last ten years have seen the development of invertebrate derived DNA (iDNA, (Calvignac-Spencer et al., 2013; Schnell et al., 2012)) and the analysis of DNA associated with environmental matrices such as soil or water (eDNA, (Harper et al., 2018; Thomsen & Willerslev, 2015)). Metabarcoding was instrumental in the emergence of these new approaches for which, by definition, there is no non-molecular alternative. Conversely, in conventional macro-organism community studies where the traditional identification using morphological criteria has been used for many years, metabarcoding is still currently marginally used as a routine identification tool. Yet, metabarcoding could enhance the capacity to characterize these communities through iDNA, eDNA or through the extraction of DNA from a homogenate of whole organisms collected together (bulk DNA; (Deiner et al., 2017)). A major limitation to the democratization of metabarcoding is the absence of standardized metabarcoding protocol that has been established and validated by the scientific community (Leese et al., 2018). In particular, methodological roadblocks are encountered throughout each step of sample processing (i.e. DNA extraction, enrichment of the targeted DNA barcode, sequencing, bioinformatic treatment and taxonomic assignment). Hereafter, we focused on two major metabarcoding roadblocks: specimen picking before extraction and DNA barcode enrichment.

Specimen picking is a major concern for bulk DNA samples and is particularly critical when the ratio of targeted organism over substrate is low (e.g. invertebrates sampled in streambed). It aims to separate individuals from substrate (e.g. leaves, sand…) prior to DNA extraction to avoid PCR inhibition (Elbrecht, Vamos, Meissner, Aroviita, & Leese, 2017) and to limit the quantity of material to be processed during DNA extraction. This step is time and cost consuming. Direct extraction from fixative agent, usually ethanol, has been proposed as a time-saver alternative to specimen picking (Hajibabaei, Spall, Shokralla, & van Konynenburg, 2012; Zizka, Leese, Peinert, & Geiger, 2018). When fixative agent is used as a DNA template, organisms are not destroyed and are conserved for further taxonomic work or downstream analyses (Leese et al., 2016). Similar techniques have already been employed by the ancient DNA community where the destruction of samples such as museum specimens is to be avoided (Paijmans, Fickel, Courtiol, Hofreiter, & Forster, 2016; van der Valk, Lona Durazo, Dalen, & Guschanski, 2017). Yet, little research on bulk DNA has been conducted on this alternative (but see Hajibabaei et al. (2012); Zizka et al. (2018)) and a rigorous comparison with traditional specimen picking is warranted before it can be used as a standard template of DNA in community studies.

PCR enrichment bias is often considered as the most problematic roadblock in metabarcoding because it may alter species detection and relative abundance recovery of species (Elbrecht & Leese, 2017; Leese et al., 2018; Piñol, Senar, & Symondson, 2018). Prior to sequencing, DNA barcodes are first amplified by PCR using primers that may not have the same number of mismatches with the targeted sequences across taxa (Piñol, Mir, Gomez-Polo, & Agustí, 2015). During a PCR, if DNA barcode sequences from two different species are in equimolar concentration, but that the first species presents less mismatches with the primers, this species’ sequence will be amplified preferentially. Consequently, amplification efficiency is expected to be non-equal across taxa, leading from under-amplification to no amplification of some taxa in the worst case scenario. PCR bias was demonstrated for fungi (Bellemain et al., 2010), bacteria (Frank et al., 2008), invertebrates (Piñol et al., 2015) and vertebrates (Arif, Khan, Al Sadoon, & Shobrak, 2011). Other factors affect PCR like GC content (Aird et al., 2011) or inhibitors that can remain after DNA extraction but primer bias is commonly presented as the major cause of biases in metabarcoding (e.g. (Elbrecht & Leese, 2015; Piñol et al., 2015; Pinto & Raskin, 2012)). In consequence, a lot of work focused on primer design to decrease PCR biases with, for instance, the use of several primer pairs, degenerated primers or amplification of several DNA barcodes (Drummond et al., 2015; Elbrecht & Leese, 2017; Elbrecht et al., 2016; Gibson et al., 2015; Jusino et al., 2019; Leray & Knowlton, 2017; Zhang, Chain, Abbott, & Cristescu, 2018). These efforts increased the species detection rate but the quantitative bias was not solved completely (Piñol et al., 2018).

Avoiding PCR enrichment will, by definition, solve the PCR bias issue (Porter & Hajibabaei, 2018). Low (Linard, Crampton-Platt, Gillett, Vogler, & Timmermans, 2015) or high (Porter & Hajibabaei, 2018) coverage metagenome sequencing (i.e. sequencing a community DNA without any enrichment) can be used to assemble entire organelle genomes. This approach provides an efficient way to recover species richness and taxa relative biomass, although the proportion of organelle reads is extremely low making metagenome sequencing much more expensive than PCR metabarcoding (Bista et al., 2018; Gómez-Rodríguez, Crampton-Platt, Timmermans, Baselga, & Vogler, 2015; Zhou et al., 2013). Furthermore, only a small part of an organelle genome is usable for taxonomic assignment as reference databases mostly contain DNA barcode sequences (e.g. COI for metazoan, 16S for bacteria, ITS for fungi, (Creer et al., 2016)). Although methods are being developed to reduce organelle genome sequencing cost (Macher, Zizka, Weigand, & Leese, 2017), the construction of exhaustive organelle genome reference databases will be a long-term and expensive process. Another PCR-free alternative is capture enrichment where targeted sequences hybridize to baits and are retrieved by magnetism (Dowle, Pochon, C. Banks, Shearer, & Wood, 2016). Contrary to metagenome sequencing, capture enrichment increases the proportion of targeted reads reducing the sequencing cost (Jones & Good, 2016). Baits are long oligonucleotides (more than 60 bp) which are designed from reference sequences. Capture enrichment has already been extensively used in genomics and genetics where it is used to retrieve thousands of loci (e.g. exons capture, (Hodges et al., 2007)) and SNPs arrays (Yang et al., 2009) of one species in a single reaction prior to sequencing. It is also commonly used for ancient DNA to both increase the amount of endogenous DNA and allow loci enrichment where DNA fragmentation tends to complicate PCR reactions (Avila-Arcos et al., 2011; Horn, 2012). Capture enrichment has also recently been used in phylogenetics where the goal is to target thousands of loci in many related species (McCormack et al., 2013; Phuong & Mahardika, 2018). Conversely, capture enrichment is not commonly used in community ecology (but see Dowle, Pochon, J, Shearer, & Wood, (2016); Shokralla et al., (2016)) which opens new challenges as in this new context the diversity of organism is extremely high but the number of targeted loci is rather small. Capture enrichment should be more robust to identify taxa in a community than PCR enrichment because (i) thousands of different baits can be designed and (ii) when sequences are unknown or species are polymorphic, few mismatches between the baits and the targeted sequences should not bias DNA enrichment as they would do with PCR primers (Li, Hofreiter, Straube, Corrigan, & Naylor, 2013; Paijmans et al., 2016; Portik, Smith, & Bi, 2016). For example, one bait designed for a species of the genus *Danio* permitted to detect others *Danio* species in the Amazon basin where the reference species was absent (Mariac et al., 2018). In a single study, capture enrichment was compared to PCR enrichment and was found to detect more taxa but was unable to estimate relative abundances (Dowle et al., 2016). Liu et al. (2016) and Wilcox et al. (2018) have shown that relative abundances can be recovered with capture enrichment if species-specific corrections were to be applied to take into account variation in the number of mitochondria copy number among species (Liu et al., 2016; Wilcox et al., 2018). Such corrections are unfortunately not suitable to the complexity of field samples. Dowle et al. (2016) study was based on natural communities only described at a coarse taxonomic level (family to genus level) and with indirect biomass measurements. Thus, the capacity of capture enrichment to describe both community diversity, and relative biomass without species-specific biomass corrections, requires further testing with controlled communities.

In this study, we first investigated the capacity of cytochrome oxidase subunit I (COI) gene capture enrichment to detect taxa and retrieve initial biomass without species specific correction. Second, we evaluated if DNA extracted from ethanol (etDNA) can be used as an alternative to organism picking and homogenization (bulk DNA) by assessing species detection and initial biomass recovery of this alternative template DNA with PCR and capture enrichment. Tests were carried out using two types of freshwater mock communities (MC): (i) low diversity MC (10 species) with variable dry biomass across taxa and (ii) high diversity MC (52 taxa) with homogeneous biomass across taxa from (Elbrecht & Leese, 2015).

## MATERIALS & METHODS

### 1. Mock community design

Freshwater macrozoobenthos specimens for the 10 species mock communities (MC) were sampled from various streams in east of France during May 2017, except for two species, *Gammarus fossarum* and *Chironomus riparius*, which came from Irstea livestock, France (ECOTOX team, RiverLy, France). The 10 species were chosen to represent a wide taxonomic range and because they can be easily identified to the species level by the naked eye (table 1). Two hundred milliliters of ethanol 96% (EtOH) were dispense in bottles and individuals were placed alive in the ethanol. Eight samples with different relative biomass for the 10 species were constructed (Figure 1, table 1). Samples were then stored for 6 months at 4°C until DNA extraction. DNA was extracted from the whole organisms (bulk DNA) and from the preservative EtOH (etDNA).

**Table 1.** Dry biomass in mg and number of individuals (italic) of freshwater invertebrate species in the 10 species mock communities.

| Species | *Mock community* |  |  |  |  |  |  |  |
| --- | --- | --- | --- | --- | --- | --- | --- | --- |
|  | 1 | 2 | 3 | 4 | 5 | 6 | 7 | 8 |
| *Chironomus riparius*<br>Meigen, 1804 | 1.84 (*5*) | 1.83 (*5*) | 2.58 (*5*) | 2.42 (*5*) | 3.66 (*10*) | 4.62 (*11*) | 3.89 (*10*) | 4.44 (*10*) |
| *Epeorus assimilis*<br>Eaton, 1885 | 102.32 (*5*) | 55.53 (*6*) | 77.89 (*5*) | 96.47 (*5*) | 62.94 (*4*) | 110.73 (*4*) | 114.80 (*4*) | 74.23 (*4*) |
| *Heptagenia sulphurea*<br>(O.F. Müller, 1776) | 15.73 (*5*) | 20.79 (*5*) | 21.08 (*5*) | 18.02 (*5*) | 38.31 (*10*) | 28.19 (*10*) | 32.58 (*10*) | 38.37 (*10*) |
| *Isoperla rivulorum*<br>(Pictet, 1841) | 37.97 (*5*) | 31.88 (*5*) | 36.28 (*5*) | 30.89 (*5*) | 35.11 (*6*) | 33.39 (*6*) | 36.22 (*6*) | 39.47 (*6*) |
| *Nemurella picteti*<br>Klapálek, 1900 | 2.83 (*5*) | 2.49 (*5*) | 2.34 (*5*) | 3.60 (*5*) | 0.50 (*1*) | 0.97 (*1*) | 0.77 (*1*) | 0.56 (*1*) |
| *Hydropsyche siltalai*<br>Doehler, 1963 | 68.42 (*5*) | 53.15 (*5*) | 53.63 (*5*) | 53.32 (*5*) | 80.36 (*8*) | 66.27 (*8*) | 86.71 (*8*) | 79.86 (*8*) |
| *Athripsodes aterrimus*<br>(Stephens, 1836) | 6.10 (*6*) | 6.68 (*5*) | 7.87 (*5*) | 6.23 (*5*) | 9.10 (*6*) | 7.96 (*6*) | 5.39 (*6*) | 7.01 (*6*) |
| *Physella acuta*<br>(Draparnaud, 1805) | 23.30 (*5*) | 24.85 (*5*) | 23.03 (*5*) | 31.30 (*5*) | 16.81 (*4*) | 15.22 (*4*) | 13.25 (*4*) | 20.68 (*4*) |
| *Ancylus fluviatilis*<br>O.F. Müller, 1774 | 13.42 (*5*) | 18.16 (*5*) | 18.40 (*5*) | 15.42 (*5*) | 20.79 (*8*) | 20.47 (*8*) | 23.21 (*8*) | 15.98 (*8*) |
| *Gammarus fossarum*<br>Koch, 1836 | 23.07 (*5*) | 20.28 (*5*) | 16.89 (*5*) | 16.72 (*5*) | 4.4 (*1*) | 4.02 (*1*) | 5.17 (*1*) | 3.76 (*1*) |

**Figure 1:**
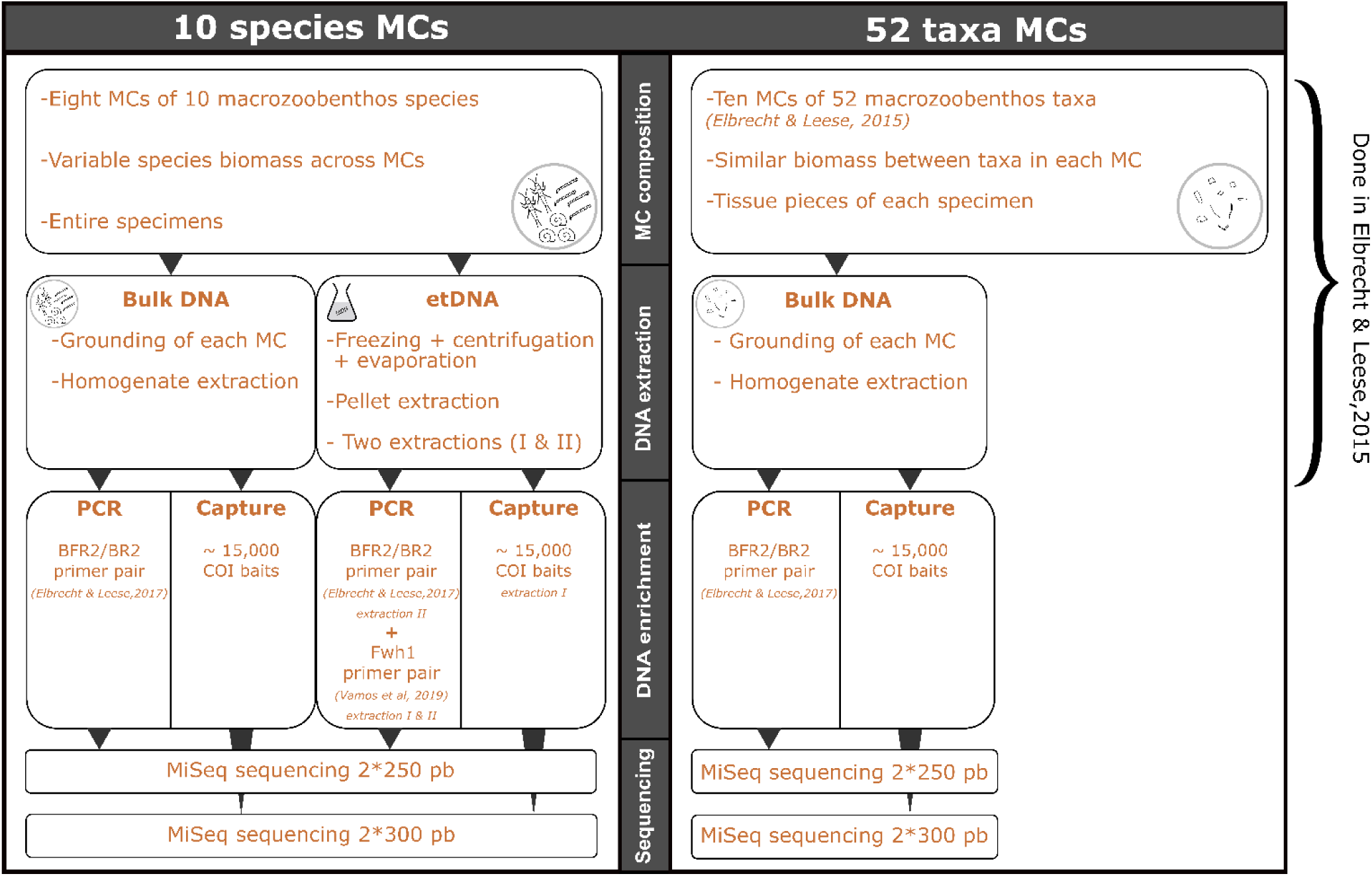
Overview of the experimental design of the study. Two different mock communities (MCs) were used: 10 species MCs and 52 taxa MCs. Ten species MCs were used to assess initial biomass recovery and etDNA performances with PCR and capture enrichment. Both MC were used to assess taxa detection. etDNA: ethanol DNA.

The 52 taxa mock communities were designed by Elbrecht and Leese (2015). In a nutshell, for ten different MC, a roughly similar dry biomass was collected from 52 taxa identified to the lowest taxonomic level based on morphology, and the homogenized tissue pool was then extracted for each sample using a salt extraction protocol (Figure 1, see Elbrecht & Leese (2015) for details). Because the biomass was fairly homogeneous among taxa in these MC, we only used these MC to compare the performance of capture and PCR enrichment in predicting species occurrences.

### 2. DNA extraction of the 10 species mock communities

#### Bulk DNA

For each MC, individuals were picked up and sorted by species in petri dishes. They were let to dry overnight and dry biomass for each species was weighted. Mollusc shells and Trichoptera cases were removed prior to the weighing. Individuals were pooled together and the entire community was grounded with a bead mill MM200 (Retsch) during 4 min at 30 Hz. The whole homogenate was extracted (mean ± SD: 245.8 ± 28.6 mg) with FastDNA® Spin Kit for Soil (MP Biomedicals, USA) following manufacturer’s protocol. The extracted DNA was purified with Agencourt AMPure XP purification beads (Beckman Coulter, USA) to further remove solvents. A negative extraction control was also made, following the same extraction protocol but without starting material.

#### etDNA

For each mock community, roughly 50 mL of preservative ethanol was collected. Glycogen and sodium acetate were added to precipitate DNA and samples were placed at -80°C for at least 72 hours. After centrifugation, ethanol was removed and after total ethanol evaporation, the dried residue was dissolved in the buffer solution of NucleoSpin Tissue® kit (Macherey-Nagel Gmbh, Germany) and DNA was extracted following manufacturer’s instructions. As the amount of DNA was limiting, extractions were repeated twice: a first extraction to compare PCR and capture enrichment methods (extraction I); a second to compare two different primer pairs performance for this supposedly degraded DNA template (extraction II). Four negative extraction controls were also made during extraction II by filling tubes with 96% ethanol which were then extracted following the same protocol.

### 3. COI mock community reference databases

The 10 species reference database was built by sequencing the COI Folmer region of each species of the 10 species mock communities (Folmer, Black, Hoeh, Lutz, & Vrijenhoek, 1994). One specimen per species was extracted following the same protocol as for bulk DNA. PCR reactions were performed in a total volume of 25 µL with 1X of PCR standard buffer (including 3 mM MgCl2, Eurobio, France), 0.05 U/µL of EurobioTaq DNA polymerase (Eurobio, France), 0.8 mM of each dNTP (Eurogentec, Belgium), 0.1 mg/mL of BSA (New England BioLabs, USA), 0.4 µM of each primer (LCO1490/HCO2198, Table S1) and 0.5 µL of template DNA. The amplification consisted in an initial denaturation at 95°C for 3 min, followed by 40 cycles of denaturation at 95°C for 20 s, annealing at 51°C for 30 s and extension at 72°C for 45 s, with a final extension at 72°C for 5 min. Purified template DNA was sequenced on both strands with the PCR primers using standard Sanger sequencing (Biofidal, France). All sequences were manually checked for errors and cleaned up with Finch TV software 1.4.0. This protocol did not work for one species (*Heptagenia sulphurea*) and its sequence was downloaded from NCBI to complete the 10 species reference database (GenBank Accession Number HE651395.1).

The 52 taxa reference database was already available for the 52 taxa MC (Elbrecht & Leese, 2017). It includes the haplotypes of all organisms used in their experiment, leading to a reference database of 212 COI sequences. For one taxon (Nematoda), Elbrecht and Leese (2017) could not get any amplification. Since this taxon is missing in Elbrecht and Leese (2017) haplotype reference database, it was not taken into account for downstream analysis.

### 4. Amplicon sequencing and analysis

#### Library preparation and sequencing

For the 10 species and 52 taxa MC bulk DNA and etDNA (extraction II), a 421 bp fragment within the COI Folmer region was amplified with the BF2/BR2 primer set with Illumina Nextera tails which is well suited for freshwater macrozoobenthos (Elbrecht & Leese, 2017), Table S1). For etDNA extractions I and II, a shorter region of 178 bp (Fwh1 primer set with Illumina Nextera tails, (Vamos, Elbrecht, & Leese, 2017), Table S1) was also targeted. PCR reactions were performed in triplicates in a total volume of 25 µL with 1X of PCR buffer (including 3mM MgCl2 and 400 μM each dNTP, QIAGEN® Multiplex PCR kit, Qiagen, Germany), 0.5 µM of each primer and 10 ng of template DNA for bulk DNA or 5 µL of template DNA for etDNA. The amplification consisted in an initial denaturation at 95°C for 5 min, followed by 25 (bulk DNA) or 35 (etDNA) cycles of denaturation at 95°C for 30 s, annealing at 50°C (bulk DNA) or 52°C (etDNA) for 30 s and extension at 72°C for 2 min, with a final extension at 72°C for 10 min. Triplicate PCR products were pooled and purified with Agencourt AMPure XP purification beads (Beckman Coulter, USA) and quantified using QuantiFluor® dsDNA System (Promega, USA). Ten ng of each purified PCR products were then used in a second PCR to dual index each sample with a unique tag combination (Table S3 for tag combinations details) and to add Illumina adapters. PCR reactions were performed in a total volume of 25 µL with 1X of PCR buffer (BIOAmp® Blend Mix, Biofidal, France), 2.5 nM of MgCL2, 200 µM of each dNTP, 0.25 µM of Illumina primer (Table S1) and 0.02 U/µL of HOT BioAmp Taq (Biofidal, France). PCR products were purified with Agencourt AMPure XP purification beads (Beckman Coulter, USA), quantified using QuantiFluor® dsDNA System (Promega, USA) and pooled at the same concentration (2 nM) for sequencing. The PCR amplicon libraries were sequenced using a 2*250 paired-end V3 MiSeq sequencing kit (Biofidal, France). Negative controls were included to every PCR during the library preparation.

#### Bioinformatic analysis

Reads were delivered demultiplexed and adapter trimmed. The reads of the 52 taxa mock communities, bulk DNA and etDNA of the 10 species mock communities were processed independently with the same bioinformatic pipeline. First, forward and reverse reads were merged with Vsearch 2.8.4 with a minimum overlapping of 10 nucleotides and a maximum difference of the overlapping region of 5 nucleotides (Rognes, Quince, Nichols, Flouri, & Mahé, 2016). Then, the primer regions were removed from the merged reads with cutadapt 1.9.1 (M. Martin, 2014) and the reads were quality filtered (maximum expected error (ee) of 1, minimum length of 200, no Ns allowed) and dereplicated with Vsearch. Sequences observed less than twice (< 2 reads) were removed. Then, chimeras were *de novo* detected and removed with Vsearch. Finally, sequences were clustered with a similarity cutoff of 97% identity into MOTUs (Molecular Operational Taxonomic Units). MOTUs were assigned to species using the 10 species reference database or the 52 taxa reference database with the blastn algorithm (Camacho et al., 2009). Only alignments with an e-value under 1E-10, a query cover over 200 bases for bulk DNA and over 90 bases for etDNA and an identity over 97% were conserved as good alignments for further analysis. MOTUs were also compared to a complete COI protein reference database made from 8 taxa (*Asellus aquaticus* ADA69754.1, *Daphnia pulex* AAD33231.1, *Dinocras cephalotes* AGZ03516.1, *Gammarus fossarum* YP_009379680.1, *Physella acuta* YP_008994230.1, *Radix balthica* HQ330989.1, *Sericostoma personatum* AJR19241.1, *Thremma gallicum* AJR19254.1) present in one or both MCs using diamond 0.9.22 (blastx, more sensitive option, e-value threshold of 1E-10 (Buchfink, Xie, & Huson, 2014)). This allowed us to detect COI sequences even if they did not belong to the species used in the MC. We also estimated the rate of contamination by comparing the quality filtered reads to a protein database containing the proteomes of 100 eukaryotic species from Ensembl (Zerbino et al., 2017) and Ensembl Metazoa (Kersey et al., 2017) as well as the proteomes of 837 prokaryotic species retrieved from the Microbial Genome Database for Comparative Analysis (Uchiyama, Mihara, Nishide, & Chiba, 2015) selecting one species per genus. Sequences were assigned to coarse taxonomic groups (archaea, eubacteria, fungi, plant, protist, protostomia – here Arthropoda, Annelida, Brachiopoda, Mollusca, Nematoda and Platyhelminthes – and other metazoa – corresponding to Deuterostomia) using the best diamond hit (blastx, more sensitive option, e-value threshold of 1E-10).

### 5. Capture sequencing and analysis

#### Bait design

While one of the primary goals of the present study is to assess capture versus PCR enrichment efficiency using mock communities, we chose to develop a larger set of baits that could be used in future assessment of French freshwater macrozoobenthos diversity. Up to now and aside from non-Chironomidae dipteran, 3,245 macrozoobenthos species belonging to 22 orders have been identified in small and medium streams of France (Aukema & Rieger, 2013; Coppa, 2019; D’Hondt & Ben Ahmed, 2009; Dusoulier, 2008; Gargominy et al, 2011; Grand & Boudot, 2007; Henry & Magniez, 1983; Le Doaré & Vinçon, 2019; Pattée & Gourbault, 1981; Piscart & Bollache, 2012; Queney, 2011; Serra et al, 2015; Souty-Grosset et al, 2006; Thomas, 2019; Tillier, 2019; Vallenduuk, 2004). One hundred twenty two species of six orders can be considered as marginal in streams and were thus discarded. All the available COI sequences were downloaded for the 3,123 remaining species from GenBank and BOLD with PrimerMiner 0.18 in December 2016 (Elbrecht & Leese, 2016). Four orders had less than half of their known taxa with available COI barcode and were discarded. Within the 12 remaining orders, the species without COI sequences were barcoded whenever possible, i.e. when tissues could be obtained. Organisms were extracted using Chelex (BioRad, USA) and the PCR and sequencing conditions were identical to those of section 3 of the methods. Sequences were deposited on GenBank (97 species, GenBank Accession Numbers MK584300:MK584515). At the end, 1,525 species out of 1,689 known species had a DNA barcode available, representing more than 90% of the targeted species (Table S2). For bait development, sequences were processed by order. First, COI sequences were aligned with blastx 2.7.1 to a reference *Drosophila yakubai* COI sequence (Accession Number: NC_001322.1) and identical sequences were collapsed using a perl script (http://github.com/TristanLefebure/collapse_to_uniq_seq). Then, 120 bp baits were constructed *in silico* using BaitFisher with a tilling of 60 bp and a cluster threshold of 5%, leading to a total of 15,038 baits generated from 100,367 unique sequences (Mayer et al., 2016). The COI *in silico* baits were then sent for RNA bait synthesis to Arbor Biosciences (USA).

#### Library preparation, hybridization and sequencing

Starting quantity of DNA was 1 µg for bulk DNA (10 and 52 taxa MC) and between 14 and 57 ng for etDNA (extraction I). DNA of each sample was sheared into approximately 600 bp nucleotide fragments by ultrasound sonication with a Qsonica Q800R (Qsonica, USA). Library preparation was conducted using NEBNext® Ultra™ II DNA Library Prep Kit for Illumina® (New England BioLabs, USA) following manufacturer’s instructions. Briefly, after sheared DNA end repair, the 5’ ends were phosphorylated and the 3’ ends were A-tailed. Then, Illumina Nextera tails (Table S1) were ligated to the DNA fragments followed by a clean-up and a size selection of 500-700 nucleotides long fragments with Agencourt AMPure XP purification beads (Beckman Coulter, USA). Finally, DNA fragments were amplified to dual index the libraries (Table S3 for tag combinations details) and to add Illumina adapters (Table S1). COI capture enrichment was conducted using myBaits® Custom kit following manufacturer’s instructions (Arbor Biosciences, USA).

One hundred ng of library DNA was used for capture enrichment for bulk DNA and 230 ng for etDNA following manufacturer’s instructions for degraded DNA. Baits were diluted 10 times and hybridization lasted 24h for both DNA templates with the exception that hybridization for etDNA was done at 55°C instead of 65°C following manufacturer’s instructions for degraded DNA. The final library amplification was performed in a total volume of 50 µL per reaction with the KAPA HiFi DNA Polymerase (Kapa Biosystems, USA) with P5 and P7 Illumina primers (Table S1) using the following conditions: initial denaturation at 98°C for 2 min, followed by 21 cycles of denaturation at 98°C for 20 s, annealing at 60°C for 30 s and extension at 72°C for 1 min, with a final extension at 72°C for 5 min. Capture library concentrations were determined by qPCR with a KAPA qPCR kit (KAPA Library Quant Kit, Kapa Biosystems, USA) and pooled at the same concentration for sequencing. The capture libraries were sequenced using a 2*300 paired-end V3 MiSeq sequencing kit (Biofidal, France).

#### Bioinformatic analysis

Reads were delivered demultiplexed and adapter trimmed. As for PCR analysis, the reads of the 52 taxa MC, bulk DNA and etDNA of the 10 species MC were processed independently with the same bioinformatic pipeline. First, forward and reverse reads were merged with Vsearch with a minimum overlapping of 10 nucleotides and a maximum difference of the overlapping region of 5 nucleotides (Rognes et al., 2016). Because shearing could lead to fragment longer than 600 nucleotides, merged and non-merged sequences were conserved for downstream bioinformatic steps. Reads were then quality filtered (maximum ee of 1, minimum length of 150, 50 Ns allowed) with Vsearch. Because the bait set contained baits for other genes for other projects (i.e. 16S, NAD1, NAD4, NAD5, CYTB, and ATP6) and because their presence in the bait set can alter downstream results, the reads corresponding to these genes were removed from the quality filtered reads in all downstream analysis. They were recovered with a blastn on a reference database (blastn-short, e-value threshold of 1E-10) containing the complete sequence of NAD1, NAD4, NAD5, CYTB and ATP6 genes of 8 species (*Asellus aquaticus* ADA69754.1, *Daphnia pulex* AAD33231.1, *Dinocras cephalotes* AGZ03516.1, *Gammarus fossarum* YP_009379680.1, *Physella acuta* YP_008994230.1, *Radix balthica* HQ330989.1, *Sericostoma personatum* AJR19241.1, *Thremma gallicum* AJR19254.1), the 16S of the 10 species MC (Table S1) and the 16S corresponding to the 52 taxa MC (downloaded from NCBI). The remaining reads were assigned to species using the 10 species reference database or the 52 taxa reference database using BLAST algorithm (Camacho et al., 2009). Only alignments with an e-value below 1E-10, a query cover over 250 and an identity over 97 were used for taxonomic assignment. As for amplicon, filtered reads were also compared to a complete COI protein reference database to estimate the total number of COI reads and to a protein database to estimate the rate and origin of contaminations.

### 6. Capture efficacy

Capture efficacy (sensus (Cha & Thilly, 1993)) was evaluated by measuring the percentage of targeted reads for each capture library (i.e. capture specificity) and the X-fold enrichment (i.e. capture efficiency) as in Maggia et al. (2017). The percentage of targeted reads is the ratio of the number of reads assigned with diamond on the COI protein database or with blastn to a 16S, NAD1, NAD4, NAD5, CYTB and ATP6 nucleotide reference database to the number of quality filtered reads. To estimate the X-fold enrichment, four samples from the 10 species MC were sequenced without any enrichment. The X-fold enrichment was calculated using the ratio of the percentage of targeted reads from the capture library to the percentage of targeted reads from the enrichment-free library.

### 7. Species detection and initial biomass recovery

To compensate for sequencing effort as well as total biomass variation among samples, read count assigned to species and biomass (mg) were transformed in proportion of reads (i.e. ratio of the number of COI reads assigned to the species to the total number of COI reads assigned to the COI mock community reference database) and proportion of biomass (i.e. ratio of the biomass of the species to the total biomass of the MC), respectively.

#### Species detection

A species was considered present in a sample when represented by at least one read for the capture sequencing pipeline and one MOTU for the amplicon sequencing pipeline. We calculated the detection sensitivity (*S*) which measures the number of detection success to the total number of trials 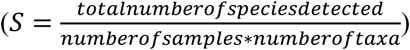

#### Biomass recovery

For the bulk DNA and etDNA of the 10 species MC, logistic models were built to investigate the relationship between read proportion and biomass proportion. For each DNA template and enrichment method, we compared sets of mixed effects models (i.e. no fixed effect and biomass as fixed effect with no random effect, random intercept, random intercept and slope by species or samples) using Aikake’s Information Criterion (AIC, (Burnham & Anderson, 2002)). We also summarized to what extent each method was able to predict species biomass by comparing the observed read proportions to the expected read proportions where read proportion perfectly predict biomass proportion (i.e. a y=x relationship). To this aim, we calculated the mean absolute error (MAE) for each method as follow: 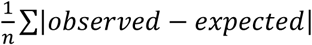 (Willmott & Matsuura, 2005). The lower the MAE is, the closer the observed read proportions are to the expected read proportions.

All statistical analyses were conducted with R (R Core team, 2018). All logistic models relating biomass proportion to read proportion were fitted using lme4 package (glmer functions, with a Logit link and a Binomial family, (Bates, Mächler, Bolker, & Walker, 2014)).

## RESULTS

### 1. Sequencing results

For bulk DNA, amplicon sequencing produced 60,337 to 365,331 raw reads and capture sequencing 165,089 to 857,040 raw reads per sample. For etDNA, amplicon sequencing produced 209,425 to 365,331 raw reads for Fwh1 primer pair of extraction I, 219,124 to 314,441 raw reads for Fwh1 primer pair of extraction II, 172,036 to 280,065 raw reads for BF2/BR2 primer pair of extraction II and capture sequencing 217,240 to 911,484 raw reads (Table 2). In both approaches, the capture sequencing effort was higher but a higher proportion of reads was discarded through the bioinformatic pipeline, particularly for etDNA (Table 2).

**Table 2.** Assessment of PCR and capture enrichment specificity on bulk and ethanol DNA. “% of targeted reads”: for capture enrichment only, percentage of reads that align to the COI, 16S, NAD1, NAD4, NAD5, CYTB, and ATP6 reference databases, “% of COI reads”: percentage of reads that align to the COI protein reference database, “% of COI reads assigned to a MC species”: percentage of the number of reads that were successfully assigned to a species using the COI nucleotide reference databases to the COI assigned reads and “% of non-protostomian reads”: percentage of reads that align to non-protostomian groups. Indicated values are mean number and standard deviation. For sample values, see Table S4 (PCR enrichment) and Table S5 (Capture enrichment). 10 sps: 10 species MC; 52 taxa:52 taxa MC; Ø means that the measure cannot be calculated.

| **bulk DNA** | *Mock community* | *Raw reads* | *Quality filtered reads* | *% targeted reads* | *% COI reads* | *% of COI reads assigned to a MC species* | *% of non-protostomian reads (contaminants)* |
| --- | --- | --- | --- | --- | --- | --- | --- |
| *PCR enrichment* | 10 sps | 170,026 ± 40,272 | 114,701 ± 31,160 | Ø | 49.54 ± 4.83 | 96.01 ± 3.24 | 2.77 ± 1.30 |
|  | 52 taxa | 135,168 ± 40,871 | 91,935 ± 32,720 | Ø | 42.33 ± 4.61 | 96.55 ± 3.80 | 2.37 ± 1.36 |
| *Capture enrichment* | 10 sps | 493,967 ± 288,240 | 243,483 ± 86,526 | 43.34 ± 24.52 | 25.16 ± 18.81 | 61.17 ± 5.55 | 10.73 ± 3.78 |
|  | 52 taxa | 553,825 ± 165,868 | 264,694 ± 75,864 | 80.21 ± 5.09 | 69.30 ± 6.51 | 63.27 ± 1.96 | 5.31 ± 1.70 |
| *Enrichment-free* | 10 sps | 921,863 ± 68,162 | 574,443 ± 43,547 | 1.19 ± 0.31 | 0.023 ± 0.028 | 26.14 ± 9.21 | 31.66 ± 1.96 |
| **etDNA** | *Mock community* | *Raw reads* | *Quality filtered reads* | *% targeted reads* | *% COI reads* | *% of COI reads assigned to a MC species* | *% of non-protostomian reads (contaminants)* |
| *PCR enrichment - Fwh1 extraction I* | 10 sps | 283,644 ± 52,944 | 233,856 ± 43,256 | Ø | 62.97 ± 4.47 | 46.36 ± 11.47 | 18.61 ± 12.78 |
| *PCR enrichment - Fwh1 extraction II* | 10 sps | 205,229 ± 81,671 | 169,772 ± 69,282 | Ø | 66.34 ± 5.88 | 47.72 ± 14.65 | 10.13 ± 4.12 |
| *PCR enrichment - BF2/BR2 extraction II* | 10 sps | 203,406 ± 43,493 | 123,148 ± 47,936 | Ø. | 48.97 ± 9.81 | 60.95 ± 18.88 | 28.21 ± 19.29 |
| *Capture Enrichment – extraction I* | 10 sps | 517,202 ± 234,942 | 309,637 ± 147,304 | 27.18 ± 8.39 | 19.09 ± 5.82 | 33.71 ± 10.96 | 56.81 ± 10.17 |

### 2. Capture efficacy

The percentage of targeted reads after capture enrichment (i.e. capture specificity) ranged from 16.24 to 86.66% (mean ± se: 63.82% ± 24.83) for bulk DNA and from 19.52 to 40.79% (mean ± se: 27.18 ± 8.39) for etDNA (Table 2, Table S5). For bulk DNA, the 52 taxa MC showed higher and more homogeneous capture specificity (mean ± se: 80.21 ± 5.09) in comparison with bulk DNA of the 10 species MC (mean 43.34 ± 24.52). The average X-fold enrichment (i.e. capture efficiency) was 974 (range: 214-2470) meaning that on average 974-fold more targeted reads were sequenced with capture enrichment than without any enrichment.

The percentage of COI reads that were assigned to a MC species was very heterogeneous among DNA template and enrichment methods. Amplicons on bulk DNA gave the best results (mean: 96.45% COI assignment), followed by capture on bulk (62.33 %), amplicon on etDNA (46.36 %) and finally capture on etDNA (33.71 %). Reads that do not match to the reference MC COI database can have multiple origins including contamination from other organisms. The majority of reads were assigned to protostomians for all but one experiment – capture enrichment on etDNA – where half of the reads belonged to eubacteria and the other half to protostomians (Table 2, Figure S2).

### 3. PCR and capture enrichment for bulk DNA

#### Species detection

Capture enrichment detected systematically more species (*S*=0.96) than PCR (*S*=0.68) among the 10 species MC (Figure 2). In PCR libraries, three species belonging to Gastropoda (*Ancylus fluviatilis* and *Physella acuta*) and Amphipoda *(Gammarus fossarum)* were never detected. With capture enrichment, every species was detected in 5 out of 8 samples (Figure 2). In the 52 taxa MC, we found the same pattern, with capture enrichment (*S*=0.96) detecting more species than PCR enrichment (*S*=0.93) (Figure 2). Each enrichment method eventually failed to detect a small but different set of taxa.

**Figure 2.**
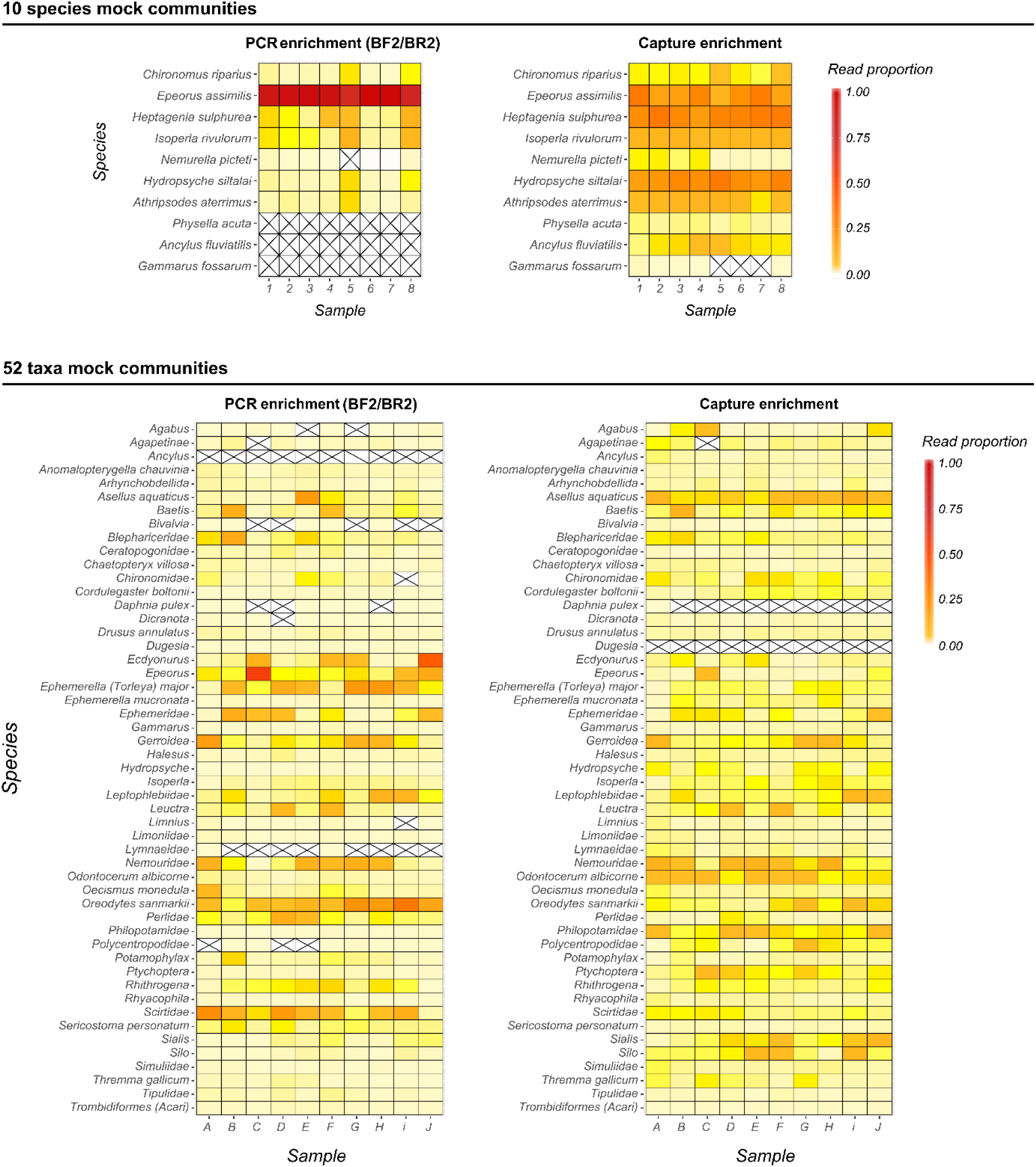
Taxa recovery performance assessed using read proportion for PCR (BF2/BR2 primer pair, left) and capture (right) enrichment and for bulk DNA of the 10 species mock communities (MCs) (top) and bulk DNA of the 52 taxa MC (bottom). Read proportions are shown for each taxa (in rows) and mock community (in column). A crossed out cell indicates no assigned read. For detailed read counts, see Tables S6 (10 species MCs) and S7 (52 taxa MCs).

#### Biomass recovery

Whatever the enrichment method, following the AIC, the best mixed model to predict initial species biomass using species read proportion was the model combining biomass as a fixed effect and species as a random effect on the intercept and slope (Table 3). This suggests that the mean read proportion between species is different independently of their biomass and that the relationship between biomass proportion and read proportion also differs among species (Figure 3). Using this type of mixed models, a significant relationship between biomass and read proportion was found for both enrichment methods (PCR: p-value=0.024; capture: p-value=0.001, Figure 3). The mean absolute errors between observed and expected read proportion if biomass proportions were to be perfectly translated into read proportions (i.e. y=x relationship) were higher for PCR (MAE=0.11) than for capture (MAE=0.056). Under a scenario where there is no relationship between biomass and read proportions and where each species contribute to the same read proportion independently of its biomass (i.e. 1/10 read proportion), the MAE would be of 0.07, again highlighting that the PCR enrichment step wiped out most of the biomass signal. While absolute biomass variations may be lost, we also tested if biomass ranks could be recovered using Spearman’s rank correlation coefficient. Again PCR enrichment performed poorly compared to capture enrichment (average Spearman rho 0.53 and 0.67 for PCR and capture enrichment, respectively).

**Table 3.**
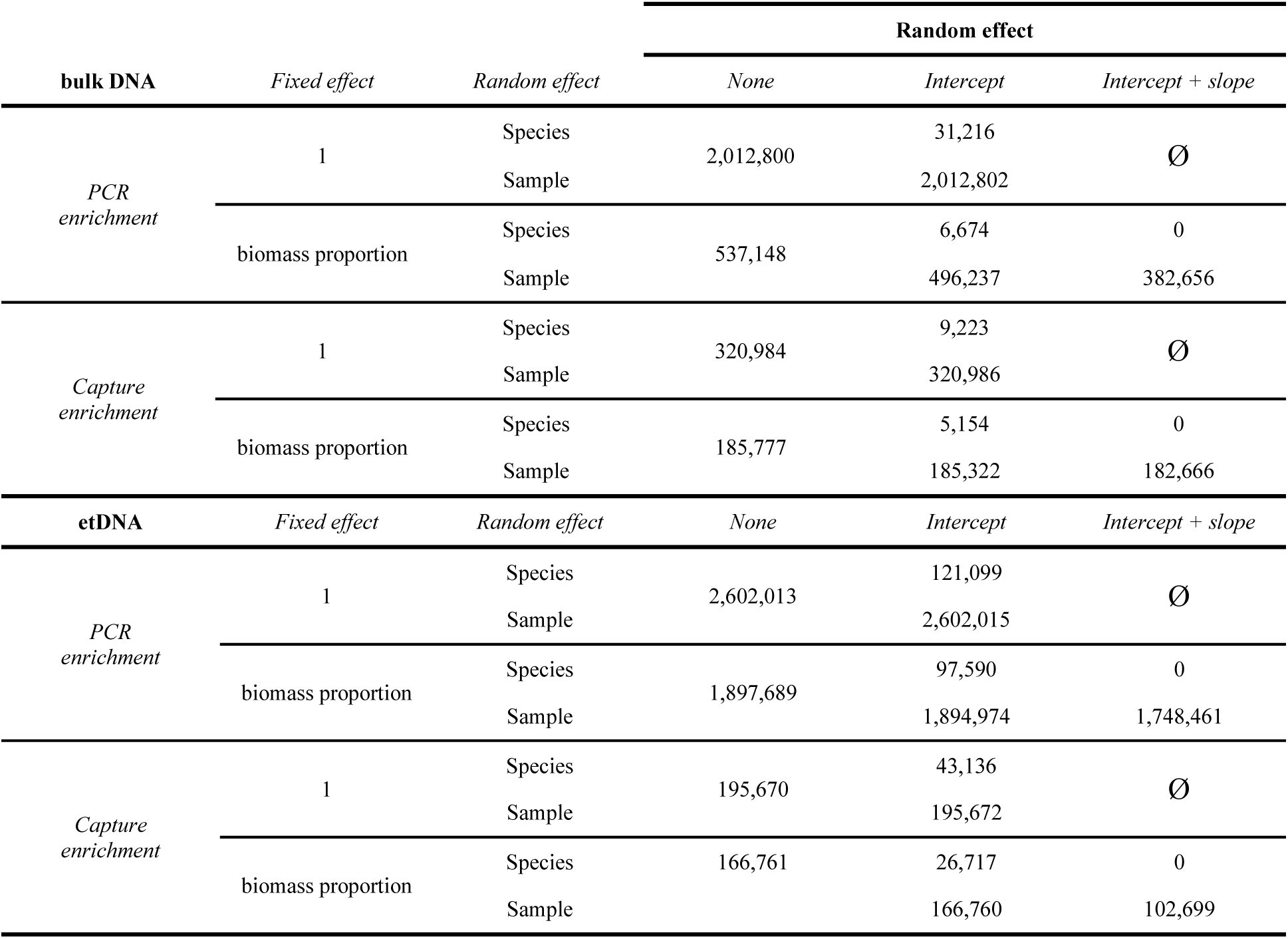
Testing the link between read proportion (dependant variable) and initial biomass proportion. Only relative ΔAIC to the best model are shown (i.e. AICmodel - AICbest model). Models with and without fixed effect (biomass proportion) and two random effects (species and sample) were built. For each random effect, three models were built: no random effect, random intercept and random intercept and slope. Best models correspond to the models with relative ΔAIC of 0. Ø means that the model could not be tested.

**Figure 3:**
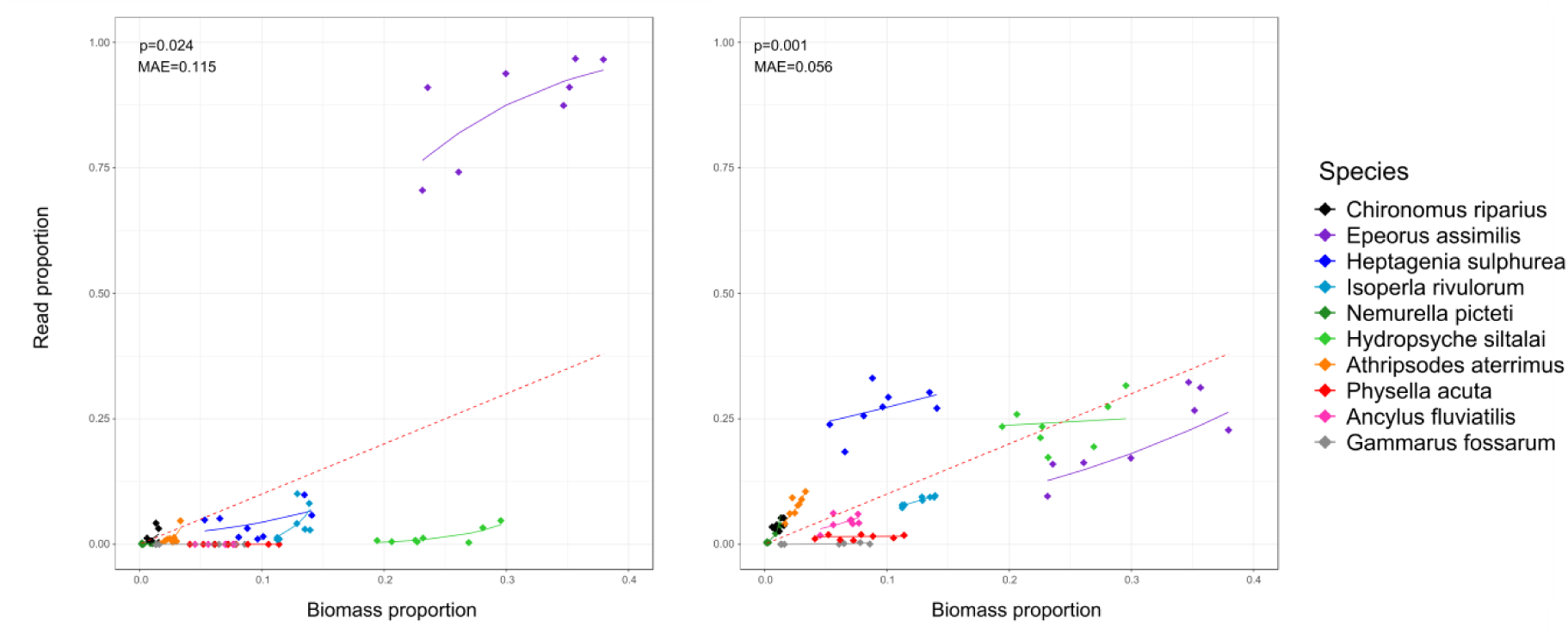
Relationship between biomass proportion and read proportion for PCR (left) and capture (right) enrichment method. Logistic mixed model predictions with random intercept and slope by species are shown (solid line). The dashed red line corresponds to the expected linear relationship where one unit of read proportion equals to one unit of biomass proportion.

### 4. etDNA performance

As for bulk DNA, capture enrichment detected more species (*S*=0.97) than PCR (*S*=0.85) (Figure 4, extraction I) on the etDNA template. Concerning PCR, the primer pair Fwh1 performs better (S=0.86) than the primer pair BF2/BR2 (S=0.7) on etDNA (Figure 4, extraction II). This was expected as the long BF2/BR2 fragment may be difficult to amplify on degraded DNA. Nevertheless the two primers pairs also failed to detect different species probably indicating a primer bias rather than a degradation problem. To confirm this, we tested if the sample composition was mostly driven by DNA template and enrichment method using a Principal Coordinate Analysis (PCoA). Samples clustered by enrichment method and PCR primers but not by DNA template (Figure 5). Therefore, in this experiment, the DNA template had little to no impact compared to the enrichment method and primers. Concerning initial biomass recovery, the same models as for bulk DNA were selected for etDNA using the AIC (Table 3). However, contrary to bulk DNA, for both enrichment methods, no significant relationship between biomass proportion and read proportion was found (PCR: p-value=0.37; capture: p-value=0.58). The difficulty to infer biomass from read proportion using etDNA was also supported by high MAE values (0.102 and 0.104 for PCR and capture enrichment, respectively) and low rank correlations (average Spearman rho= 0.40 and 0.28 for PCR and capture enrichment, respectively).

**Figure 4:**
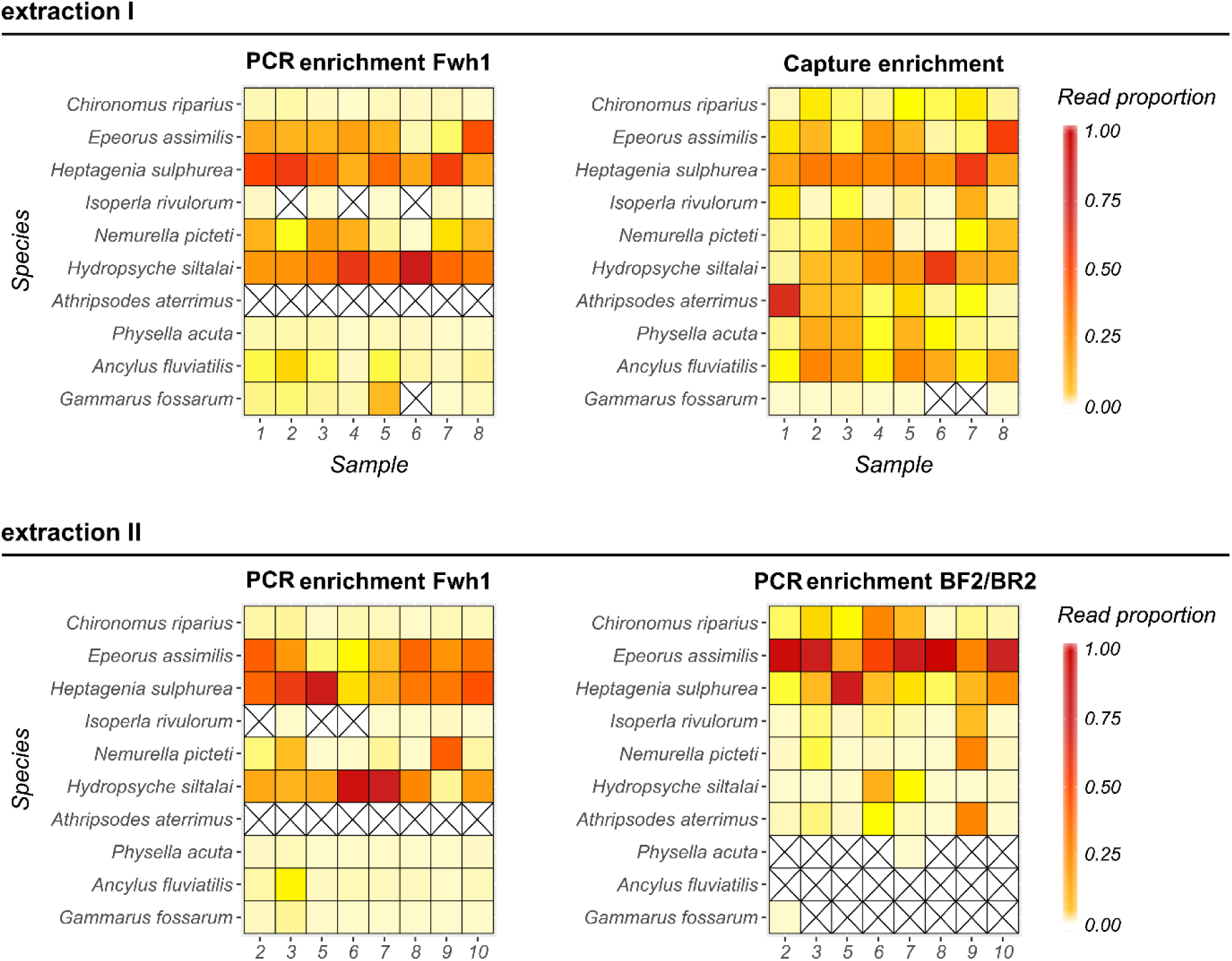
Taxa recovery performance assessed using read proportion for etDNA of the 10 species mock communities. Differences between PCR (top left) and capture (top right) enrichment were assessed using a first ethanol extraction (extraction I). Differences between Fwh1 (bottom left) and BF2/BR2 (bottom right) primer pairs were assessed using a second ethanol extraction (extraction II). Sequence abundances are shown for each taxa (in rows) and mock community (in column). A crossed out cell indicates no assigned read. For detailed read counts, see Tables S6.

**Figure 5:**
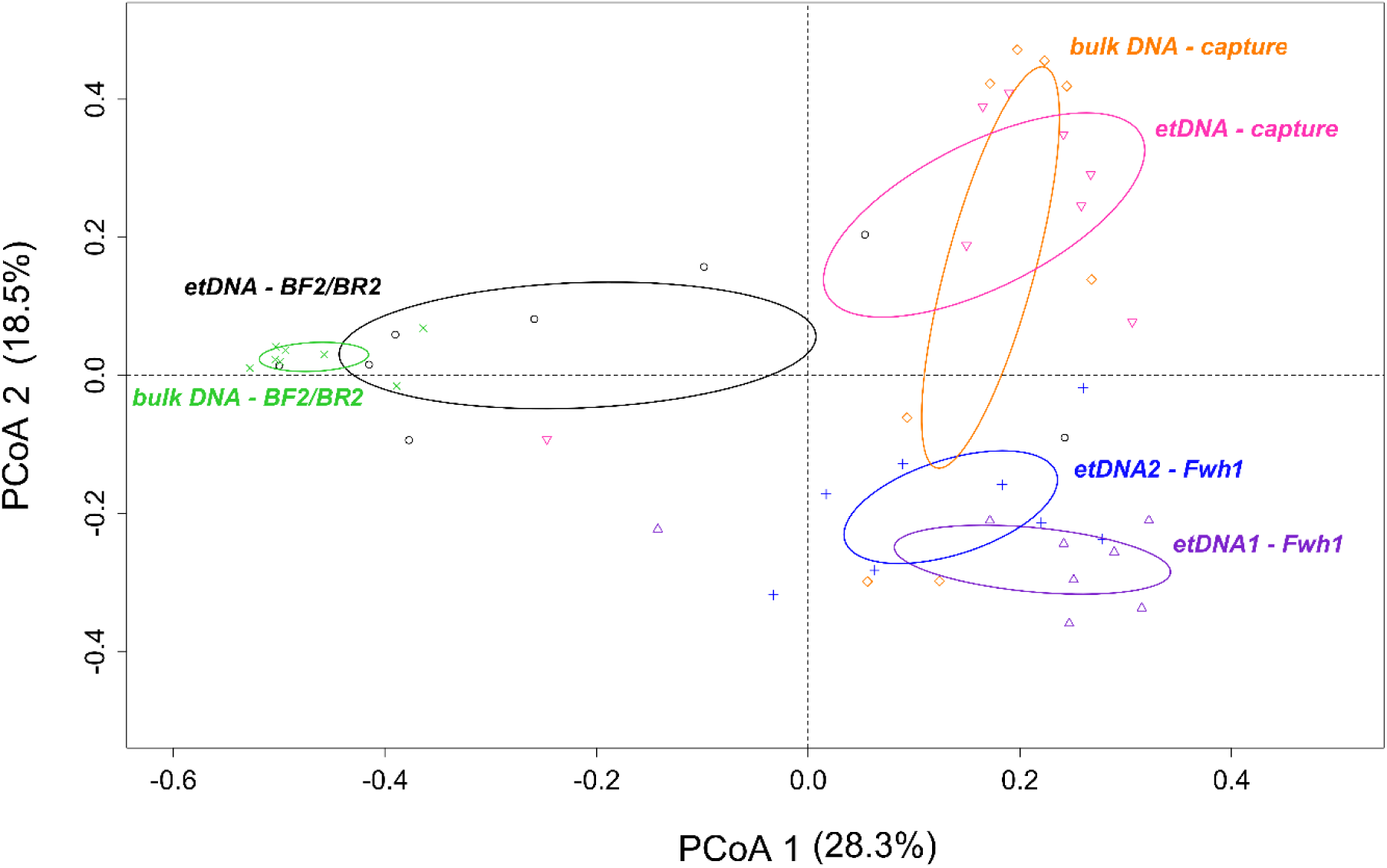
Principal Coordinate Analysis based on dissimilarity (Jaccard distances) in read proportions of each pair of the 10 species mock community samples. Samples are classified following their DNA template (etDNA and bulk DNA) and enrichment methods (BF2/BR2 amplicon, Fwh1 amplicon and capture). etDNA1 and etDNA2 correspond to the first and second extraction of DNA from ethanol, respectively. Percentage of variance explained by each axis is shown in bracket. Ellipses are drawn with a confidence limit of 0.95. Samples are grouped by enrichment method rather than by DNA template.

## DISCUSSION

### Capture versus PCR enrichment for metabarcoding

Capture enrichment consistently led to higher species detection rates compared to PCR enrichment whatever the DNA template or sample taxonomic complexity. Using capture in the 10 species MC, only one species was missed in one third of the samples where it was at low abundance (dry biomass < 1 mg). In the 52 taxa MC, capture enrichment also globally performed better that PCR enrichment in terms of species detection (20 against 35 failed detections, respectively). Both methods also missed different sets of taxa. In PCR enrichment, the commonly missed taxa systematically presented 1 or 2 mismatches with the primer pair (Figure S1) reinforcing the view that primer mismatches are the main cause for species non-detection (Elbrecht et al., 2017). Regarding capture enrichment, 8 taxa had no bait designed for them (i.e. Ceratopogonidae, Blephariceridae, Dicranota, Simuliidae, Tipulidae, Trombidiformes, Dugesia and *Daphnia pulex*) but only two taxa (*Daphnia pulex* and Dugesia) were almost systematically missed. These two taxa have COI haplotypes that are at a divergence of at least 15% to any bait whereas the 6 detected taxa have haplotypes that are closer to baits originally designed for other taxa (Figure S1). The divergence to the oligonucleotide has already been described has an important parameter for species detection (Liu et al, 2015; Portik et al, 2016; Vallender et al, 2011; van der Valk et al, 2017). In particular, Liu et al (2015) using a pool of 49 taxa found a lower enrichment efficiency when the divergence to the bait was higher than 20%. Portik et al (2016), this time working on a multi-loci capture data set, found a negative linear relationship between the probability of sequencing a locus and the divergence to the bait. This translated into a sharp decline from 60% to 20% chance of sequencing when the divergence increased from 10% to 20%. Altogether, for both enrichment methods, the divergence to the oligonucleotide (i.e. primer or bait) appears to be a determinant factor for species detection. Nevertheless, capture enrichment is much more robust to this divergence issue as it combines three characteristics that PCR enrichment misses. First, capture enrichment is less sensitive to mismatches (18 mismatches or 15% divergence) than PCR (2 mismatches or 5% divergence). Second, because thousands of different baits are designed for a given 120-base region, the probability to encounter mismatches is reduced with capture. Finally, several 120-base regions (5 in this study leading to a total cover of 600 bases) are targeted, increasing the probability to design at least one functional bait for a given species even if this species was not in the database used to design the baits. Alternatively species detection can be improved in traditional PCR-based metabarcoding by targeting multiple DNA fragment (e.g. (Drummond et al., 2015; Zhang, Chain, Abbott, & Cristescu, 2018)). This is also likely to be true but easier to implement with capture enrichment where baits for different loci can be used simultaneously (e.g. Liu et al, 2016). The ability to detect more robustly known and unknown biodiversity using a single enrichment step is a pivotal step towards a better understanding of ecosystem function and structure.

Capture enrichment also yields read proportions that are better predictors of species relative biomasses than PCR enrichment for bulk DNA. Both enrichment methods presented a positive relationship between relative biomass proportion and read proportion but this relationship was closer to a linear y=x relationship for capture enrichment. That said, a random species effect still remained in the capture enrichment model. Its origin could be attributed to a bait bias similar to the primer bias encountered in PCR enrichment but it can also be explained by other factors. Indeed, using a mitochondrial marker to reconstruct an entire dry biomass community is eventually doomed to fail. First, the amount of mitochondrial DNA belonging to a species in a sample will vary as a function of the average number of mitochondrial genomes per cell which is likely to vary extensively among species, life stages and tissues (Rooney et al, 2015). Wilcox et al. (2018) found that total initial species DNA abundances could be recovered using capture enrichment if a correction was applied to normalize for variation in the number of mitochondrial genome per cell. Second, dry biomass proportions are a very rough estimator of cell numbers. Thus, variability in cell number per biomass and in mitochondria per cell combines to blur the biomass/read proportion relationship. A similar mock community experiment but with precise knowledge of the number of mitochondrial genome per species in a sample is needed to better test for the existence of bait biases. One can argue that even if capture enrichment yields good estimates of each species mitochondrial genome proportion in a sample, this estimate will still be a poor biomass or abundance proxy. Interestingly, here we found that read proportions from gene capture enrichment already provides biomass rank estimates which in many ecological contexts, for instance in bioassessment (Elbrecht et al., 2017), would already be informative and may be sufficient.

### Ethanol DNA as a fast alternative to bulk DNA for species detection

In agreement with Martins et al. (2019), our results demonstrate the potential of etDNA to replace bulk DNA for macrozoobenthos samples if only species occurrence data is required. Species detection was not determined by DNA template but by the enrichment method and the primer pair used. Hence, species detection with etDNA was equivalent to species detection with bulk DNA. Nonetheless, no relationship between read proportion and biomass proportion was highlighted for etDNA. For biomass recovery to work with etDNA, each species should release a quantity of DNA in the ethanol proportionally to its biomass. However, the release of DNA in the ethanol might differ among development stages or species. When put alive in ethanol, some individuals regurgitate and release a lot of their DNA (Anderson et al., 2013). For example, in the class Gastropoda, one sub-class has an operculum which isolates them from the environment. When put in ethanol, these gastropods close their shell and the quantity of DNA released will be lower than for other gastropods without operculum. Also, after ethanol fixation, depending on the presence of a shell or a thick exoskeleton, the amount of released DNA during the soaking period may also be extremely variable across taxa. In conclusion, while DNA from ethanol offers a fast and non-destructive way to identify the species in a sample, differences in the way species release their DNA in ethanol seem to prevent the use of ethanol for quantitative estimates. Compared to bulk DNA, DNA extracted from preservative ethanol is more fragmented and much less concentrated. In addition to environmental contaminants such as the bacterial contaminants observed in this study, this template may also be more susceptible to reagent contamination and cross-contamination. The adoption of this alternative template may therefore need stricter sampling, sample handling and DNA extraction protocols than the one regularly used in community ecology laboratories.

### Optimising capture enrichment for metabarcoding

Despite capture enrichment already delivers better results than PCR enrichment in terms of species detection or biomass prediction, the efficiency and specificity of this emerging approach can still be optimized for metabarcoding purposes. Although capture enrichment was efficient to enrich the COI sequences of the macrozoobenthos present in our samples with an enrichment of at least 214-fold compared to non-enriched libraries, the percentage of targeted reads was highly heterogeneous and low for some samples. The specificity of the capture enrichment was particularly low when the diversity of the sample was low (43% of targeted reads for the 10 species MC compared to 80% for the 52 taxa MC). How the taxonomic diversity of a sample influences the efficiency of capture enrichment remains unexplained and warrants further experiments where the impact of a gradually increasing sample diversity is tested while controlling for other factors such as the extraction protocol or the overall phylogenetic diversity.

The specificity of capture enrichment estimated by the number of COI reads that matched a taxa included in the MC was systematically lower than with PCR enrichment (about 40% of the COI reads did not match a MC taxa). When looking at the coarse taxonomic composition of the COI reads sequenced after capture enrichment, we found that for bulk DNA, most of the reads (> 95%) were assigned to protostomians ruling out the possibility that bacterial or plant COI contaminations alone are reducing capture specificity. DNA from other protostomians that interact with the taxa included in the MC (e.g. preys or parasites) may contaminate the samples and reduce capture specificity. Nevertheless, we would expect the same to happen with PCR enrichment which is not the case. Alternatively, Li, Schroeder, Ko, and Stoneking (2012) found that the robustness of capture to mismatches is also a drawback when targeting mitochondrial genes: in addition to capture the targeted mitochondrial gene, it also captures nuclear copies of this gene (NUMTs). The number of NUMT loci per genome is highly variable across taxa but can easily reach hundreds to thousands of loci (Hazkani-Covo, Zeller, & Martin, 2010). In some species there might therefore be more NUMTs in a cell than there are mitochondrial genomes. Given that baits can capture divergent loci and that there is many different baits, NUMTs may represent a significant burden in mitochondrial capture enrichment. Finally, capture enrichment also increase the sequencing coverage of the flanking regions of the targeted sequences (Portik et al, 2016). With the pipeline used in this study, the reads containing more than 41% of a flanking region will not be classified as a read coming from a targeted region. The lower specificity of capture enrichment observed in our samples, but not observed with PCR enrichment, might therefore be explained by a combined effect of all these factors.

For capture enrichment on etDNA, we estimated that almost half of the reads belonged to eubacteria and the other half to protostomians. This is likely to be linked to the regurgitation of gastric microbiota and the release of epidermic microbiota by some species. By this process, bacterial DNA may actually dominate the ethanol DNA. In this experiment, a lower hybridization temperature was used for etDNA (55°C) than for bulk DNA (65°C) in the hope of capturing more fragmented DNA. Retrospectively, this lower temperature is likely to have decreased the specificity of the capture enrichment and increased the representation of bacterial DNA. Indeed, the specificity of capture enrichment is higher for degraded DNA and samples with a lot of contaminants when a higher hybridization temperature is used (Paijmans et al., 2016). Noteworthy, 95% of the bacterial reads did not share any similarity with the mitochondrial COI, again reinforcing the idea that the large amount of bacterial reads in etDNA samples is probably the result of the combination of non-stringent hybridization conditions and bacterial DNA dominance in etDNA.

As found in this study and others (e.g. Liu et al., 2016; Mariac et al., 2018; Portik et al, 2015), the divergence between the targeted DNA and the baits is a critical factor in capture enrichment. As such, to recover all taxa in a community, the bait design must be optimized to reduce this divergence. The priority is thus to design baits using a reference database that is exhaustive. While reference DNA barcodes are available for most species for some groups, this is far from being the case for many groups (e.g. (Sonet et al., 2013)). Knowing the divergence threshold of 15% (this study) over which species are not captured and detected, alternative strategies could be deployed to barcode key missing taxa in the reference databases. Indeed, a very limited set of baits is sufficient to represent a whole family or a genus if the intra-group genetic diversity is lower than the divergence threshold (Mariac et al., 2018). This diversity is heterogeneous between taxa group, for example the maximum intra-family COI divergence in Gammaridea (Amphipoda) is 30% but is only 17% in Chloroperlidae (Plecoptera). Therefore, prior investigations are needed to establish the diversity of each group and the number of taxa that have to be described before a robust set of baits can be designed. Another alternative would be to design baits from a set of representative COI sequences and mutate them according to a given divergence to obtain a set of baits that can hybridize to most sequences. Such baits would permit to capture non-barcoded or even new species but could also reduce capture specificity by capturing untargeted sequences such as bacterial genes or NUMTs.

In addition to the bait design strategy, other capture enrichment parameters may need further optimization for a metabarcoding application. The hybridization temperature is known to be a major factor controlling capture efficacy (Li et al., 2013). However, Paijmans et al. (2016) compared different hybridization temperatures but unexpectedly did not find any interaction between hybridization temperature and distance to the bait. But Paijmans et al. (2016) used a pool of 21 feline species, with all the probes designed using a single reference species. Therefore their results probably need to be tested on a data-set that is more similar to a metabarcoding data-set, with more divergent taxa and probes designed from a large database, before the interaction between hybridization temperature and distance to the bait can be completely dismissed. The number of pooled libraries in a single hybridization reaction is another potentially important parameter. While pooling many samples in a single capture reaction significantly cuts the hybridization budget, Portik et al. (2016) showed that it significantly reduces the complexity of the sequencing libraries. In a metabarcoding context, this may result in a lower species discovery rate in disfavor of the rare species. In the same vein, the number of capture rounds is a substantial parameter increasing capture efficacy but also the hybridization budget (Li et al., 2013; Mariac et al., 2018; Templeton et al., 2013; van der Valk et al., 2017). With a second round of capture, Mariac et al. (2018) increased capture specificity from 0.57% to 70.5% and capture efficiency by 40 times but with the penalty of doubling hybridization costs. After sequencing capture enriched metabarcoding libraries, another challenge is the bioinformatic treatment of the reads. Indeed, processing capture data targeting one or few DNA barcodes for many species is considerably different than assembling multiple targeted genes for a single organism. To our knowledge, while there is several assembly pipelines that were developed for the latter case (co-assembly of different loci from a single species, (e.g. (Allio et al., 2019)), there is no available bioinformatic pipelines for the former. Assembling metabarcoding capture reads is challenging as it requires the independent assembly of orthologous sequences belonging to different species which may be more or less divergent and more or less polymorphic. An alternative to assembly is to directly map the capture reads on a reference database as done in this study. While functional, this solution is sub-optimal because capture reads do not start and end at the same position as in amplicon sequencing. This complicates the read clustering and taxonomic assignment, with many capture reads probably lost as they only partially map to the reference library. While the sequencing of the flanking regions of targeted region would probably be highly valuable for taxonomic assignment, as long as there is no dedicated bioinformatic solution, it paradoxically complicates the processing and analysis of capture reads. Finally, a comparison of these mock community results with field samples in which the diversity is much higher and where the template is also much more complex is warranted to confirm the potential of capture enrichment in community ecology.

## CONCLUSION

Capture enrichment is a promising alternative to PCR enrichment for metabarcoding. Its main advantage is to provide better species detection thanks to its robustness to mismatches. At this point, while performing much better than PCR enrichment, absolute biomass reconstruction is not applicable without species mitogenome copy number correction and is therefore not tractable for community studies where hundreds of species can be encountered. Yet, biomass rank ordination appears to be robust and could be used for ecological purpose. Albeit more bacterial contamination, the use of etDNA coupled to capture enrichment presents interesting compromises. It saves a lot of time by ignoring the organism picking step and permits to save organisms for further analyses. However, etDNA should be used for studies where quantitative information is not required. Finally, these promising results call for additional testing of the bait design strategy and hybridization reaction parameters, as well as testing capture enrichment on complex field samples.

## ACKNOLEDGEMENTS

We thank Maxence Forcellini, Bertrand Launay and Guillaume Le Goff for morphological macrozoobenthos identification and fieldwork assistance. We thank Clémentine François and Florian Leese for advices and assistance in bioinformatic analysis and bait design. We thank the three anonymous reviewers for their helpful and relevant comments on the manuscript. This study was supported by the French Biodiversity Agency (AFB) “Grant 26: headwater biodiversity dynamics: a molecular perspective”, and the CNRS Mission pour les Initiatives Transverses et Interdisciplinaires (project XLIFE CAPTAS).

## DATA ACCESSIBILITY

Additional reference sequences developed for bait design are available on Genbank (Accession Numbers MK584300:MK584515). COI and 16S sequences barcoded for the 10 species MC assignment are available on GenBank (Accession Numbers MK584516:MK584524 for COI and MK584525:MK584534 for 16S). The bait set designed for this study is available on Zenodo (Zenodo (https://doi.org/10.5281/zenodo.2581410). FASTQ files from PCR and capture enrichment are available on NCBI (Accession Number SRA SRP188737). Scripts for bioinformatic analysis are available on GitHub (https://github.com/mailysgauthier/bioinf-cap-PCR).

## AUTHOR CONTRIBUTIONS

Samples of the 10 species MC were collected by MG. Methodology design was conceived by MG, TL, CD, VE and TD. Laboratory work was conducted by MG, LKD, AN. Data analysis was conducted by MG, TL and VE. MG and TL led the writing of the manuscript. All authors contributed to write the manuscript.

## REFERENCES

Abdelfattah, A., Malacrinò, A., Wisniewski, M., Cacciola, S. O., & Schena, L. (2018). Metabarcoding: A powerful tool to investigate microbial communities and shape future plant protection strategies. Biological Control, 120, 1–10. doi:10.1016/j.biocontrol.2017.07.009

Aird, D., Ross, M. G., Chen, W. S., Danielsson, M., Fennell, T., Russ, C., … Gnirke, A. (2011). Analyzing and minimizing PCR amplification bias in Illumina sequencing libraries. Genome Biology, 12, R18. doi:10.1186/gb-2011-12-2-r18

Allio, R., Schomaker-Bastos, A., Romiguier, J., Prosdocimi, F., Nabholz, B., & Delsuc, F. (2019). doi:10.1101/685412

Anderson, J. T., Zilli, F. L., Montalto, L., Marchese, M. R., McKinney, M., & Park, Y.-L. (2013). Sampling and Processing Aquatic and Terrestrial Invertebrates in Wetlands. In Wetland Techniques (Vol. 1, pp. 143–195).

Arif, I. A., Khan, H. A., Al Sadoon, M., & Shobrak, M. (2011). Limited efficiency of universal mini-barcode primers for DNA amplification from desert reptiles, birds and mammals. Genetics and Molecular Research, 10, 3559–3564. doi:10.4238/2011.October.31.3

Aukema, B. & Rieger C. (1995) Catalogue of the Heteroptera of the Palaeartic Region. Vol. 1: Enicocephalomorpha, Dipsocoromorpha, Nepomorpha, Gerromorpha and Leptopodomorpha. Netherlands Entomological Society, Wageningen, The Netherlands. 222 p.

Aukema, B., Rieger, C. & Rabitsch, W. (2013) Catalogue of the Heteroptera of the Palaearctic Region. Vol. 6: Supplement. Netherlands Entomological Society, Wageningen, The Netherlands. 629 p.

Avila-Arcos, M. C., Cappellini, E., Romero-Navarro, J. A., Wales, N., Moreno-Mayar, J. V., Rasmussen, M., … Gilbert, M. T. (2011). Application and comparison of large-scale solution-based DNA capture-enrichment methods on ancient DNA. Sci Rep, 1, 74. doi:10.1038/srep00074

Baird, D. J., & Hajibabaei, M. (2012). Biomonitoring 2.0: a new paradigm in ecosystem assessment made possible by next-generation DNA sequencing. Molecular Ecology, 21, 2039–2044. doi:10.1111/j.1365-294X.2012.05519.x

Bates, D., Mächler, M., Bolker, B., & Walker, S. (2014). Fitting Linear Mixed-Effects Models using lme4. doi:10.18637/jss.v067.i01

Bellemain, E., Carlsen, T., Brochmann, C., Coissac, E., Taberlet, P., & Kauserud, H. (2010). ITS as an environmental DNA barcode for fungi: An in silico approach reveals potential PCR biases. BMC Microbiology, 10, 1–9. doi:10.1186/1471-2180-10-189

Bista, I., Walsh, K., Zhou, X., Seymour, M., Christmas, M., Bradley, D., … Creer, S. (2018). Performance of amplicon and shotgun sequencing for accurate biomass estimation in invertebrate community samples. Molecular Ecology Resources, 18, 1020–1034. doi:10.1111/1755-0998.12888

Bortolus, A. (2008). Error Cascades in the Biological Sciences: The Unwanted Consequences of Using Bad Taxonomy in Ecology. AMBIO: A Journal of the Human Environment, 37, 114–118. doi:10.1579/0044-7447(2008)37[114:ecitbs]2.0.co;2

Bringloe, T. T., Cottenie, K., Martin, G. K., & Adamowicz, S. J. (2016). The importance of taxonomic resolution for additive beta diversity as revealed through DNA barcoding. Genome, 1140, 1130–1140.

Buchfink, B., Xie, C., & Huson, D. H. (2014). Fast and sensitive protein alignment using DIAMOND. Nature Methods, 12, 59–60. doi:10.1038/nmeth.3176

Burnham, K. P., & Anderson, D. R. (2002). Model Selection and Multimodel Inference: A Practical Information-Theoretic Approach (2nd ed). Ecological Modelling, 172, 488. doi:10.1016/j.ecolmodel.2003.11.004

Calvignac-Spencer, S., Merkel, K., Kutzner, N., Kuhl, H., Boesch, C., Kappeler, P. M., … Leendertz, F. H. (2013). Carrion fly-derived DNA as a tool for comprehensive and cost-effective assessment of mammalian biodiversity. Mol Ecol, 22(4), 915–924. doi:10.1111/mec.12183

Camacho, C., Madden, T. L., Ma, N., Coulouris, G., Avagyan, V., Bealer, K., & Papadopoulos, J. (2009). BLAST+: architecture and applications. BMC Bioinformatics, 10, 421. doi:10.1186/1471-2105-10-421

Cha, R. S., & Thilly, W. G. (1993). Specificity, efficiency, and fidelity of PCR. Genome Res., 3, 18–29.

Creer, S., Deiner, K., Frey, S., Porazinska, D., Taberlet, P., Thomas, K., … Bik, H. (2016). The ecologist’s field guide to sequence-based identification of biodiversity. Methods in Ecology and Evolution, 56, 68–74. doi:10.1111/2041-210X.12574

Deiner, K., Bik, H. M., Machler, E., Seymour, M., Lacoursiere-Roussel, A., Altermatt, F., … Bernatchez, L. (2017). Environmental DNA metabarcoding: Transforming how we survey animal and plant communities. Mol Ecol, 26(21), 5872–5895. doi:10.1111/mec.14350

de Moraes-Barros, N., Silva, J. A. B., & Morgante, J. S. (2011). Morphology, molecular phylogeny, and taxonomic inconsistencies in the study of Bradypus sloths (Pilosa: Bradypodidae) Journal of Mammalogy, 92, 86–100. doi:10.1644/10-mamm-a-086.1

D’Hondt, J. L., & BEN A., R. (2009). Catalogue et clés tabulaires de détermination des Hirudinées d’eau douce de la faune Française. Bulletin de la Société zoologique de France, 134, 263–298.

Dowle, E. J., Pochon, X., C. Banks, J., Shearer, K., & Wood, S. A. (2016). Targeted gene enrichment and high-throughput sequencing for environmental biomonitoring: a case study using freshwater macroinvertebrates. Molecular Ecology Resources, 16, 1240–1254. doi:10.1111/1755-0998.12488

Drummond, A. J., Newcomb, R. D., Buckley, T. R., Xie, D., Dopheide, A., Potter, B. C. M., … Nelson, N. (2015). Evaluating a multigene environmental DNA approach for biodiversity assessment. GigaScience, 4. doi:10.1186/s13742-015-0086-1

Elbrecht, V., & Leese, F. (2015). Can DNA-based ecosystem assessments quantify species abundance? Testing primer bias and biomass-sequence relationships with an innovative metabarcoding protocol. PLoS ONE, 10, 1–16. doi:10.1371/journal.pone.0130324

Elbrecht, V., & Leese, F. (2016). Development and validation of DNA metabarcoding COI primers for aquatic invertebrates using the R package “PrimerMiner …. PeerJ, 1–23. doi:10.7287/peerj.preprints.2044v1

Elbrecht, V., & Leese, F. (2017). Validation and Development of COI Metabarcoding Primers for Freshwater Macroinvertebrate Bioassessment. Frontiers in Environmental Science, 5, 1–11. doi:10.3389/fenvs.2017.00011

Elbrecht, V., Taberlet, P., Dejean, T., Valentini, A., Usseglio-Polatera, P., Beisel, J.-N., … Leese, F. (2016). Testing the potential of a ribosomal 16S marker for DNA metabarcoding of insects. PeerJ, 4, e1966. doi:10.7717/peerj.1966

Elbrecht, V., Vamos, E. E., Meissner, K., Aroviita, J., & Leese, F. (2017). Assessing strengths and weaknesses of DNA metabarcoding-based macroinvertebrate identification for routine stream monitoring. Methods in Ecology and Evolution, 8, 1265–1275. doi:10.1111/2041-210X.12789

Folmer, O., Black, M., Hoeh, W., Lutz, R., & Vrijenhoek, R. (1994). DNA primers for amplification of mitochondrial cytochrome c oxidase subunit I from diverse metazoan invertebrates. Molecular Marine Biology and Biotechnology, 3(5), 294–299.

Forshaw, J. M. (2010). Parrots of the World (P. U. Press Ed. Vol. 70).

Frank, J. A., Wilson, B. A., Weisbaum, J. S., Sharma, S., Reich, C. I., & Olsen, G. J. (2008). Critical Evaluation of Two Primers Commonly Used for Amplification of Bacterial 16S rRNA Genes. Applied and Environmental Microbiology, 74, 2461–2470. doi:10.1128/aem.02272-07

Forster, D. W., Bull, J. K., Lenz, D., Autenrieth, M., Paijmans, J. L. A., Kraus, R. H. S., … Fickel, J. (2018). Targeted resequencing of coding DNA sequences for SNP discovery in nonmodel species. Mol Ecol Resour, 18(6), 1356–1373. doi:10.1111/1755-0998.12924

Gargominy, O., Prié, V., Bichain, J. M., Cucherat, X., & Fontaine, B. (2011). Liste de référence annotée des mollusques continentaux de France. MalaCo, 7, 307–382.

Gibson, J. F., Shokralla, S., Curry, C., Baird, D. J., Monk, W. A., King, I., & Hajibabaei, M. (2015). Large-scale biomonitoring of remote and threatened ecosystems via high-throughput sequencing. PLoS ONE, 10, e0138432. doi:10.1371/journal.pone.0138432

Gómez-Rodríguez, C., Crampton-Platt, A., Timmermans, M. J. T. N., Baselga, A., & Vogler, A. P. (2015). Validating the power of mitochondrial metagenomics for community ecology and phylogenetics of complex assemblages. Methods in Ecology and Evolution, 6, 883–894. doi:10.1111/2041-210X.12376

Grand, D., & Boudot, J. P. (2007). Les libellules de France, Belgique et Luxembourg. Biotope

Hajibabaei, M., Baird, D. J., Fahner, N. A., Beiko, R., & Golding, G. B. (2016). A new way to contemplate Darwin’s tangled bank: how DNA barcodes are reconnecting biodiversity science and biomonitoring. Philosophical Transactions of the Royal Society of London B: Biological Sciences, 371, 20150330. doi:10.1098/rstb.2015.0330

Hajibabaei, M., Spall, J. L., Shokralla, S., & van Konynenburg, S. (2012). Assessing biodiversity of a freshwater benthic macroinvertebrate community through non-destructive environmental barcoding of DNA from preservative ethanol. BMC Ecology, 12, 28. doi:10.1186/1472-6785-12-28

Harper, L. R., Buxton, A. S., Rees, H. C., Bruce, K., Brys, R., Halfmaerten, D., … Hänfling, B. (2018). Prospects and challenges of environmental DNA (eDNA) monitoring in freshwater ponds. Hydrobiologia, 826(1), 25–41. doi:10.1007/s10750-018-3750-5

Hazkani-Covo, E., Zeller, R. M., & Martin, W. (2010). Molecular poltergeists: mitochondrial DNA copies (numts) in sequenced nuclear genomes. PLoS Genet, 6(2), e1000834. doi:10.1371/journal.pgen.1000834

Hebert, P. D. N., Ratnasingham, S., & de Waard, J. R. (2006). Barcoding animal life: cytochrome c oxidase subunit 1 divergences among closely related species Proceedings of the Royal Society of London. Series B: Biological Sciences, 270, 96–99. doi:10.1098/rsbl.2003.0025

Henry, J. P., & Magniez, G. (1983). Introduction pratique à la systématique des organismes des eaux continentales françaises-4. Crustacés Isopodes (principalement Asellotes). Publications de la Société Linnéenne de Lyon. 52, 319–357.

Hodges, E., Xuan, Z., Balija, V., Kramer, M., Molla, M. N., Smith, S. W., … McCombie, W. R. (2007). Genome-wide in situ exon capture for selective resequencing. Nature Genetics, 39, 1522. doi:10.1038/ng.2007.42

Horn, S. (2012). Target enrichment via DNA hybridization capture. Methods Mol Biol, 840, 177–188. doi:10.1007/978-1-61779-516-9_21

Hubert, N., Delrieu-Trottin, E., Irisson, J. O., Meyer, C., & Planes, S. (2010). Identifying coral reef fish larvae through DNA barcoding: A test case with the families Acanthuridae and Holocentridae. Molecular Phylogenetics and Evolution, 55, 1195–1203. doi:10.1016/j.ympev.2010.02.023

Ji, Y., Ashton, L., Pedley, S. M., Edwards, D. P., Tang, Y., Nakamura, A., … Yu, D. W. (2013). Reliable, verifiable and efficient monitoring of biodiversity via metabarcoding. Ecology Letters, 16, 1245–1257. doi:10.1111/ele.12162

Jones, M. R., & Good, J. M. (2016). Targeted capture in evolutionary and ecological genomics. Mol Ecol, 25(1), 185–202. doi:10.1111/mec.13304

Jusino, M. A., Banik, M. T., Palmer, J. M., Wray, A. K., Xiao, L., Pelton, E., … Lindner, D. L. (2019). An improved method for utilizing high-throughput amplicon sequencing to determine the diets of insectivorous animals. Molecular Ecology Resources, 19, 176–190. doi:10.1111/1755-0998.12951

Kersey, P. J., Stein, J., Zadissia, A., Yates, A., Paulini, M., Urban, M., … Bolser, D. M. (2017). Ensembl Genomes 2018: an integrated omics infrastructure for non-vertebrate species. Nucleic Acids Research, 46, D802–D808. doi:10.1093/nar/gkx1011

Leese, F., Altermatt, F., Bouchez, A., Ekrem, T., Hering, D., Meissner, K., … Zimmermann, J. (2016). DNAqua-Net: Developing new genetic tools for bioassessment and monitoring of aquatic ecosystems in Europe. Research Ideas and Outcomes, 2, e11321. doi:10.3897/rio.2.e11321

Leese, F., Bouchez, A., Abarenkov, K., Altermatt, F., Borja, Á., Bruce, K., … Weigand, A. M. (2018). Why We Need Sustainable Networks Bridging Countries, Disciplines, Cultures and Generations for Aquatic Biomonitoring 2.0: A Perspective Derived From the DNAqua-Net COST Action. Advances in Ecological Research, 58, 63–99. doi:10.1016/bs.aecr.2018.01.001

Leray, M., & Knowlton, N. (2017). Random sampling causes the low reproducibility of rare eukaryotic OTUs in Illumina COI metabarcoding. PeerJ, 5, e3006. doi:10.7717/peerj.3006

Leys, M., Keller, I., Räsänen, K., Gattolliat, J. L., & Robinson, C. T. (2016). Distribution and population genetic variation of cryptic species of the Alpine mayfly Baetis alpinus (Ephemeroptera: Baetidae) in the Central Alps. BMC Evolutionary Biology, 16, 1–15. doi:10.1186/s12862-016-0643-y

Li, M., Schroeder, R., Ko, A., & Stoneking, M. (2012). Fidelity of capture-enrichment for mtDNA genome sequencing: Influence of NUMTs. Nucleic Acids Research, 40. doi:10.1093/nar/gks499

Li, C., Hofreiter, M., Straube, N., Corrigan, S., & Naylor, G. J. (2013). Capturing protein-coding genes across highly divergent species. Biotechniques, 54(6), 321–326. doi:10.2144/000114039

Linard, B., Crampton-Platt, A., Gillett, C. P. D. T., Vogler, A. P., & Timmermans, M. J. T. N. (2015). Metagenome Skimming of Insect Specimen Pools: Potential for Comparative Genomics. Genome Biology and Evolution, 7, 1474–1489. doi:10.1093/gbe/evv086

Liu, S., Wang, X., Xie, L., Tan, M., Li, Z., Su, X., … Zhou, X. (2016). Mitochondrial capture enriches mito-DNA 100 fold, enabling PCR-free mitogenomics biodiversity analysis. Molecular Ecology Resources, 16, 470–479. doi:10.1111/1755-0998.12472

Macher, J. N., Zizka, V. M. A., Weigand, A. M., & Leese, F. (2017). A simple centrifugation protocol for metagenomic studies increases mitochondrial DNA yield by two orders of magnitude. Methods in Ecology and Evolution, 9(4), 1070–1074. doi:10.1111/2041-210X.12937

Maggia, M. E., Vigouroux, Y., Renno, J. F., Duponchelle, F., Desmarais, E., & Nunez, J. (2017). DNA Metabarcoding of Amazonian Ichthyoplankton Swarms. 1–14. doi:10.1371/journal.pone.0170009

Mariac, C., Vigouroux, Y., Duponchelle, F., García-Dávila, C., Nunez, J., Desmarais, E., & Renno, J. F. (2018). Metabarcoding by capture using a single COI probe (MCSP) to identify and quantify fish species in ichthyoplankton swarms. PLoS ONE, 13, 1–15. doi:10.1371/journal.pone.0202976

Martin, G. K., Adamowicz, S. J., & Cottenie, K. (2016). Taxonomic resolution based on DNA barcoding a ff ects environmental signal in metacommunity structure. Freshwater Science, 35, 701–711. doi:10.1086/686260.

Martin, M. (2014). Cutadapt removes adapter sequences from high-throughput sequencing reads. EMBnet.journal, 17, 10. doi:10.14806/ej.17.1.200

Martins, F. M. S., Galhardo, M., Filipe, A. F., Teixeira, A., Pinheiro, P., Pauperio, J., Beja, P. (2019). Have the cake and eat it: Optimizing nondestructive DNA metabarcoding of macroinvertebrate samples for freshwater biomonitoring. Mol Ecol Resour, 19(4), 863–876. doi:10.1111/1755-0998.13012

Mayer, C., Sann, M., Donath, A., Meixner, M., Podsiadlowski, L., Peters, R. S., … Niehuis, O. (2016). BaitFisher: A Software Package for Multispecies Target DNA Enrichment Probe Design. Molecular biology and evolution, 33, 1875–1886. doi:10.1093/molbev/msw056

McCormack, J. E., Harvey, M. G., Faircloth, B. C., Crawford, N. G., Glenn, T. C., & Brumfield, R. T. (2013). A phylogeny of birds based on over 1,500 loci collected by target enrichment and highthroughput sequencing. PLoS One, 8(1), e54848. doi:10.1371/journal.pone.0054848

Oksanen, J., Blanchet G.F., Friendly M., Kindt R., Legendre P., McGlinn D., Minchin P.R., O’Hara R. B., Simpson G. L., Solymos P., Henry M., Stevens H., Szoecs E. and Wagner H. (2019). vegan: Community Ecology Package. R package version 2.5-5. https://CRAN.R-project.org/package=vegan

Paijmans, J. L., Fickel, J., Courtiol, A., Hofreiter, M., & Forster, D. W. (2016). Impact of enrichment conditions on cross-species capture of fresh and degraded DNA. Mol Ecol Resour, 16(1), 42–55. doi:10.1111/1755-0998.12420

Pattée, E., & Gourbault, N. (1981). Introduction pratique à la systématique des organismes des eaux continentales françaises. 1 Turbellariés Triclades Paludicoles (planaires d’eau douce). Publications de la Société Linnéenne de Lyon, 50, 279–304.

Phuong, M. A., & Mahardika, G. N. (2018). Targeted Sequencing of Venom Genes from Cone Snail Genomes Improves Understanding of Conotoxin Molecular Evolution. Mol Biol Evol, 35(5), 1210–1224. doi:10.1093/molbev/msy034

Piñol, J., Mir, G., Gomez-Polo, P., & Agustí, N. (2015). Universal and blocking primer mismatches limit the use of high-throughput DNA sequencing for the quantitative metabarcoding of arthropods. Molecular Ecology Resources, 15, 819–830. doi:10.1111/1755-0998.12355

Piñol, J., Senar, M. A., & Symondson, W. O. C. (2018). The choice of universal primers and the characteristics of the species mixture determine when DNA metabarcoding can be quantitative. Molecular Ecology. doi:10.1111/mec.14776

Pinto, A. J., & Raskin, L. (2012). PCR biases distort bacterial and archaeal community structure in pyrosequencing datasets. PLoS ONE, 7. doi:10.1371/journal.pone.0043093

Piscart, C., & Bollache, L. (2012). Crustacés amphipodes de surface: gammares d’eau douce. Association française de limnologie. p 122.

Porter, T. M., & Hajibabaei, M. (2018). Scaling up: A guide to high-throughput genomic approaches for biodiversity analysis. Molecular Ecology, 27, 313–338. doi:10.1111/mec.14478

Prié, V., Puillandre, N., & Bouchet, P. (2013). Bad taxonomy can kill: molecular reevaluation of Unio mancus Lamarck, 1819 (Bivalvia: Unionidae) and its accepted subspecies. Knowledge and Management of Aquatic Ecosystems, 11. doi:10.1051/kmae/2013071

Queney, P. (2004). Liste taxonomique des Coléoptères ‘aquatiques’ de la faune de France (avec leur répartition sommaire). Le Coléoptériste, 7, 3–27

R Core Team (2018). R: A Language and Environment for Statistical Computing. R Foundation for Statistical Computing, Vienna.

Rognes, T., Quince, C., Nichols, B., Flouri, T., & Mahé, F. (2016). VSEARCH: a versatile open source tool for metagenomics. PeerJ, 4, e2584. doi:10.7717/peerj.2584

Schnell, I. B., Thomsen, P. F., Wilkinson, N., Rasmussen, M., Jensen, L. R., Willerslev, E., … Gilbert, M. T. (2012). Screening mammal biodiversity using DNA from leeches. Curr Biol, 22(8), R262–263. doi:10.1016/j.cub.2012.02.058

Shokralla, S., F. Gibson, J., King, I., J. Baird, D., H. Janzen, D., Hallwachs, W., & Hajibabaei, M. (2016). Environmental DNA Barcode Sequence Capture: Targeted, PCR-free Sequence Capture for Biodiversity Analysis from Bulk Environmental Samples. doi:10.1101/087437

Sonet, G., Jordaens, K., Braet, Y., Bourguignon, L., Dupont, E., Backeljau, T., … Desmyter, S. (2013). Utility of GenBank and the Barcode of Life Data Systems (BOLD) for the identification of forensically important Diptera from Belgium and France. ZooKeys, 365, 307–328. doi:10.3897/zookeys.365.6027

Souty-Grosset, C., Holdich, D., Noel, P., Reynolds, J. D., & Haffner, P. (2006). Atlas of crayfish in Europe. Muséum national d’Histoire naturelle. p188

Stein, E. D., White, B. P., Mazor, R. D., Miller, P. E., & Pilgrim, E. M. (2013). Evaluating Ethanol-based Sample Preservation to Facilitate Use of DNA Barcoding in Routine Freshwater Biomonitoring Programs Using Benthic Macroinvertebrates. PLoS ONE, 8, 1–7. doi:10.1371/journal.pone.0051273

Sweeney, B. W., Battle, J. M., Jackson, J. K., & Dapkey, T. (2011). Can DNA barcodes of stream macroinvertebrates improve descriptions of community structure and water quality? Journal of the North American Benthological Society, 30, 195–216. doi:10.1899/10-016.1

Tachet, H., Richoux, P., Bournard, M., & Usseglio-Polatera, P. (2010). Invertébrés d’eau douce: systématique, biologie, écologie. Paris: CNRS éditions.

Templeton, J. E. L. P.M. B., Llamas, B. J. S., Haak, W., Cooper, A., & Austin, J. J. (2013). DNA capture and next-generation sequencing can recover whole mitochondrial genomes from highly degraded samples for human identification. Investigative Genetics, 5(26).

Thomsen, P. F., & Willerslev, E. (2015). Environmental DNA – An emerging tool in conservation for monitoring past and present biodiversity. Biological Conservation, 183, 4–18. doi:10.1016/j.biocon.2014.11.019

Uchiyama, I., Mihara, M., Nishide, H., & Chiba, H. (2015). MBGD update 2015: Microbial genome database for flexible ortholog analysis utilizing a diverse set of genomic data. Nucleic Acids Research, 43, D270–D276. doi:10.1093/nar/gku1152

Valentini, A., Taberlet, P., Miaud, C., Civade, R., Herder, J., Thomsen, P. F., … Dejean, T. (2015). Next-generation monitoring of aquatic biodiversity using environmental DNA metabarcoding. Molecular Ecology, n/a-n/a. doi:10.1111/mec.13428

Vallenduuk, H. J., & Cuppen, M. J. (2004). The aquatic living caterpillars (Lepidoptera: Pyraloidea: Crambidae) of Central Europe. A key to the larvae and autecology. Lauterbornia, 49, 1–17.

Vamos, E., Elbrecht, V., & Leese, F. (2017). Short COI markers for freshwater macroinvertebrate metabarcoding. Metabarcoding and Metagenomics, 1, e14625. doi:10.3897/mbmg.1.14625

Van Bortel, W., Harbach, R. E., Trung, H. D., Roelants, P., Backeljau, T., & Coosemans, M. (2001). Confirmation of Anopheles varuna in Vietnam, previously misidentified and mistargeted as the malaria vector Anopheles minimus. American Journal of Tropical Medicine and Hygiene, 65, 729–732. doi:10.4269/ajtmh.2001.65.729

van der Valk, T., Lona Durazo, F., Dalen, L., & Guschanski, K. (2017). Whole mitochondrial genome capture from faecal samples and museum-preserved specimens. Mol Ecol Resour, 17(6), e111–e121. doi:10.1111/1755-0998.12699

Wilcox, T. M., Piggott, M. P., Young, M. K., McKelvey, K. S., Schwartz, M. K., & Zarn, K. E. (2018). Capture enrichment of aquatic environmental DNA: A first proof of concept. Molecular Ecology Resources, 18, 1392–1401. doi:10.1111/1755-0998.12928

Willmott, C. J., & Matsuura, K. (2005). Advantages of the mean absolute error (MAE) over the root mean square error (RMSE) in assessing average model performance. Climate Research, 30, 79–82. doi:10.3354/cr00799

Yang, H., Ding, Y., Hutchins, L. N., Szatkiewicz, J., Bell, T. A., Paigen, B. J., … Churchill, G. A. (2009). A customized and versatile high-density genotyping array for the mouse. Nature Methods, 6, 663. doi:10.1038/nmeth.1359

Zerbino, D. R., Flicek, P., Juettemann, T., Zadissa, A., Lavidas, I., Achuthan, P., … Loveland, J. E. (2017). Ensembl 2018. Nucleic Acids Research, 46, D754–D761. doi:10.1093/nar/gkx1098

Zhang, G. K., Chain, F. J. J., Abbott, C. L., & Cristescu, M. E. (2018). Metabarcoding using multiplexed markers increases species detection in complex zooplankton communities. Evolutionary Applications, 11, 1901–1914. doi:10.1111/eva.12694

Zhou, X., Li, Y., Liu, S., Yang, Q., Su, X., Zhou, L., … Huang, Q. (2013). Ultra-deep sequencing enables high-fidelity recovery of biodiversity for bulk arthropod samples without PCR amplification. GigaScience, 2, 4. doi:10.1186/2047-217X-2-4

Zizka, V. M. A., Leese, F., Peinert, B., & Geiger, M. F. (2018). DNA metabarcoding from sample fixative as a quick and voucher-preserving biodiversity assessment method. Genome, 1–41. doi:10.1139/gen-2018-0048

