## Supplementary material for "Enhancing DNA metabarcoding performance and applicability with bait capture enrichment and DNA from conservative ethanol"

Table S1: Oligonucleotides used for barcoding, PCR and capture enrichment. The modified primers for amplicon enrichment correspond to Illumina tails plus the primer from the literature for the first PCR. The tags used for dual indexing correspond to a unique combination of eight bases that are represented here as *XXXXXXXX* (Table S3).

| **Barcoding** | | |
|---|---|---|
| Primer name | Oligonuclotide sequence (5' → 3') | Literature |
| LCO1490 | GGTCAACAAATCATAAAGATATTGG | Folmer et al, 1994 |
| HCO2198 | TAAACTTCAGGGTGACCAAAAAATCA | Folmer et al, 1994 |
| 16SarDr | CGCCTGTTTAACAAAAACAT | Palumbi et al, 1991 |
| 16Sbr | CCGGTCTGAACTCAGATCACGT | Palumbi et al, 1996 |
| **PCR enrichment (first PCR)** | | |
| Primer name | Oligonuclotide sequence (5' → 3') | Literature |
| Modified primer BF2 with Illumina Nextera tail | TCGTCGGCAGCGTCAGATGTGTATAAGAGACA GGCHCCHGAYATRGCHTTYCC | Elbrecht et al, 2017 |
| Modified primer BR2 with Illumina Nextera tail | GTCTCGTGGGCTCGGAGATGTGTATAAGAGAC AGTCDGGRTGNCCRAARAAYCA | Elbrecht et al, 2017 |
| Modified primer fwhF1 with Illumina Nextera tail | TCGTCGGCAGCGTCAGATGTGTATAAGAGACA GYTCHACWAAYCAYAARGAYATYGG | Vamos et al, 2017 |
| Modified primer fwhR1 with Illumina Nextera tail | GTCTCGTGGGCTCGGAGATGTGTATAAGAGAC AGARTCARTTWCCRAAHCCHCC | Vamos et al, 2017 |
| **Capture enrichment** | | |
| | Oligonuclotide sequence (5' → 3') | Literature |
| Illumina Nextera tail P5 | TCGTCGGCAGCGTCAGATGTGTATAAGAGACA G*T | |
| Illumina Nextera tail P7 | [Phos]CTGTCTCTTATACACATCTCCGAGCCCAC GAGAC | |
| P5 | AATGATACGGCGACCACCGA | |
| P7 | CAAGCAGAAGACGGCATACGA | |
| **Indexing (both PCR (second PCR) and capture enrichment)** | | |
| | Oligonuclotide sequence (5' → 3') | Literature |
| Illumina adapter P5 + tag | AATGATACGGCGACCACCGAACAC*XXXXXXXX* TCGTCGGCAGCGTC | |
| Illumina adapter P7 + tag | CAAGCAGAAGACGGCATACGAGAT*XXXXXXXX* GTCTCGTGGGCTCGG | |

Table S2: Retained order for bait design after selection based on sequence availability and marginality in streams upon a French species list. For each order, the numbers of species with and without a barcode available for bait design are provided as well as the percentage of species with a barcode.

| Order | Number of species with a barcode | Number of species with no barcode | Percentage of species with a barcode |
|---|---|---|---|
| Amphipoda | 14 | 5 | 73.68 |
| Arhynchobdellida | 6 | 2 | 75.00 |
| Coleoptera | 346 | 70 | 83.17 |
| Diptera (Chironomidae) | 670 | 0 | 100.00 |
| Ephemeroptera | 74 | 18 | 80.43 |
| Gastropoda | 31 | 4 | 88.57 |
| Isopoda | 4 | 1 | 80.00 |
| Odonata | 23 | 7 | 76.67 |
| Plecoptera | 99 | 19 | 83.90 |
| Rhynchobdellida | 11 | 4 | 73.33 |
| Trichoptera | 232 | 34 | 87.22 |
| Venerida | 15 | 0 | 100.00 |

Table S3: Tag combination of each sample for all enrichment methods (BF2/BR2 PCR, fwh1 PCR, capture and no enrichment) and DNA templates (bulk and ethanol DNA). etDNA1 and etDNA2 correspond to the first and second extraction of DNA from ethanol, respectively.

| PCR enrichment | | | |
|---|---|---|---|
| | | **Tag i5** | **Tag i7** |
| **Bulk DNA BF2/BR2** | 1 | CTATTAAG | AGGCAGAA |
| | 2 | TTCTAGCT | TAAGGCGA |
| | 3 | GAGCCTTA | CGTACTAG |
| | 4 | CCTAGAGT | TAAGGCGA |
| | 5 | GCGTAAGA | TAAGGCGA |
| | 6 | CTATTAAG | TAAGGCGA |
| | 7 | AAGGCTAT | TAAGGCGA |
| | 8 | AAGGCTAT | CGTACTAG |
| | A | TCGACTAG | CGTACTAG |
| | B | TTCTAGCT | CGTACTAG |
| | C | CCTAGAGT | CGTACTAG |
| | D | GCGTAAGA | CGTACTAG |
| | E | CTATTAAG | CGTACTAG |
| | F | TTATGCGA | CGTACTAG |
| | G | TCGACTAG | AGGCAGAA |
| | H | TTCTAGCT | AGGCAGAA |
| | i | CCTAGAGT | AGGCAGAA |
| | J | GCGTAAGA | AGGCAGAA |
| **etDNA1 fwh1** | 1 | CTATTAAG | TCCTGAGC |
| | 2 | TCGACTAG | TCCTGAGC |
| | 3 | GCGTAAGA | TCCTGAGC |
| | 4 | CCTAGAGT | TCCTGAGC |
| | 5 | AAGGCTAT | TCCTGAGC |
| | 6 | TTCTAGCT | TCCTGAGC |
| | 7 | GAGCCTTA | TCCTGAGC |
| | 8 | TTATGCGA | TCCTGAGC |
| **etDNA2 fwh1** | 2 | CCTAGAGT | TAGGCATG |
| | 3 | TCGACTAG | TAGGCATG |
| | 5 | TTCTAGCT | TAGGCATG |
| | 6 | AAGGCTAT | TAGGCATG |
| | 7 | GAGCCTTA | TAGGCATG |
| | 8 | TTATGCGA | TAGGCATG |
| | 9 | GCGTAAGA | TAGGCATG |
| | 10 | CTATTAAG | TAGGCATG |
| **etDNA2 BF2/BR2** | 2 | CCTAGAGT | CGAGGCTG |
| | 3 | TCGACTAG | CGAGGCTG |
| | 5 | TTCTAGCT | CGAGGCTG |
| | 6 | TCGACTAG | AAGAGGCA |
| | 7 | TTCTAGCT | AAGAGGCA |
| | 8 | CCTAGAGT | AAGAGGCA |
| | 9 | GCGTAAGA | CGAGGCTG |
| | 10 | CTATTAAG | CGAGGCTG |

| Capture enrichment | | | |
|---|---|---|---|
| | | **Tag i5** | **Tag i7** |
| **Bulk DNA** | 1 | TCGACTAG | GTAGAGGA |
| | 2 | TTCTAGCT | GTAGAGGA |
| | 3 | GCGTAAGA | AAGAGGCA |
| | 4 | CTATTAAG | AAGAGGCA |
| | 5 | CCTAGAGT | GTAGAGGA |
| | 6 | GCGTAAGA | GTAGAGGA |
| | 7 | AAGGCTAT | AAGAGGCA |
| | 8 | GAGCCTTA | AAGAGGCA |
| | A | TTATGCGA | GTAGAGGA |
| | B | TCGACTAG | AAGAGGCA |
| | C | TTATGCGA | AAGAGGCA |
| | D | CCTAGAGT | CGAGGCTG |
| | E | TTCTAGCT | AAGAGGCA |
| | F | CCTAGAGT | AAGAGGCA |
| | G | GCGTAAGA | CGAGGCTG |
| | H | CTATTAAG | CGAGGCTG |
| | i | AAGGCTAT | CGAGGCTG |
| | J | GAGCCTTA | CGAGGCTG |
| **etDNA** | 1 | CTATTAAG | TAGGCATG |
| | 2 | TCGACTAG | TAGGCATG |
| | 3 | GCGTAAGA | TAGGCATG |
| | 4 | CCTAGAGT | TAGGCATG |
| | 5 | AAGGCTAT | TAGGCATG |
| | 6 | GAGCCTTA | TAGGCATG |
| | 7 | GCGTAAGA | CTCTCTAC |
| | 8 | TCGACTAG | CTCTCTAC |
| **Enrichment-free** | 2 | TTCTAGCT | CGAGGCTG |
| | 4 | TTATGCGA | CTCTCTAC |
| | 5 | GAGCCTTA | CTCTCTAC |
| | 8 | AAGGCTAT | CTCTCTAC |

Table S4: Assessment of COI PCR enrichment specificity on two types of DNA templates. "% of COI reads": percentage of reads that align to the COI protein reference database, "% of COI reads assigned to a MC species": percentage of the number of reads that were successfully assigned to a species using the COI nucleotide reference databases to the COI assigned reads and "% of non-protostomian reads": percentage of reads that align to non-protostomian groups. Indicated values are mean number and standard deviation. For sample values, see Table S4 (PCR enrichment) and Table S5 (Capture enrichment). 10 sps: 10 species MC; 52 taxa:52 taxa MC.

**Bulk DNA**

| **Samples** | MC type | Raw | Quality filtered | Dereplicated | % reads assigned to COI | % of COI reads assigned to a MC species | % of reads assigned to non-protostomian |
|---|---|---|---|---|---|---|---|
| 1 | 10 sps | 232,590 | 154,023 | 112,841 | 45.60 | 97.62 | 2.89 |
| 2 | 10 sps | 146,043 | 101,579 | 75,380 | 47.71 | 98.35 | 1.97 |
| 3 | 10 sps | 128,848 | 68,221 | 48,296 | 56.33 | 98.88 | 0.89 |
| 4 | 10 sps | 153,984 | 109,616 | 83,361 | 42.65 | 97.44 | 3.00 |
| 5 | 10 sps | 146,348 | 95,113 | 71,087 | 45.33 | 94.38 | 3.01 |
| 6 | 10 sps | 170,792 | 124,516 | 88,660 | 53.66 | 98.37 | 1.77 |
| 7 | 10 sps | 149,057 | 102,314 | 72,570 | 53.08 | 93.28 | 5.27 |
| 8 | 10 sps | 232,553 | 162,224 | 111,594 | 51.50 | 89.74 | 3.42 |
| A | 52 taxa | 125,332 | 87,765 | 63,465 | 45.29 | 93.03 | 5.83 |
| B | 52 taxa | 110,528 | 71,808 | 52,415 | 40.71 | 97.15 | 2.56 |
| C | 52 taxa | 132,420 | 93,280 | 66,136 | 51.60 | 94.30 | 1.18 |
| D | 52 taxa | 60,337 | 31,948 | 24,348 | 36.20 | 104.42 | 2.00 |
| E | 52 taxa | 106,747 | 64,778 | 46,054 | 46.14 | 98.17 | 1.18 |
| F | 52 taxa | 216,804 | 156,491 | 115,405 | 42.73 | 95.28 | 2.70 |
| G | 52 taxa | 158,951 | 109,633 | 82,051 | 40.15 | 97.97 | 2.49 |
| H | 52 taxa | 134,076 | 92,950 | 69,692 | 39.06 | 98.85 | 2.76 |
| i | 52 taxa | 157,139 | 109,428 | 79,209 | 43.91 | 95.99 | 1.63 |
| J | 52 taxa | 149,350 | 101,269 | 76,552 | 37.49 | 90.32 | 1.38 |

**etDNA**

| **Samples** | MC type | Raw | Quality filtered | Dereplicated | % reads assigned to COI | % of COI reads assigned to a MC species | % of reads assigned to non-protostomian |
|---|---|---|---|---|---|---|---|
| 1 | 10 sps | 350,270 | 285,167 | 179,267 | 61.64 | 34.66 | 19.90 |
| 2 | 10 sps | 249,675 | 205,449 | 131,510 | 63.49 | 32.57 | 46.08 |
| 3 | 10 sps | 261,439 | 216,397 | 142,800 | 58.02 | 41.80 | 18.87 |
| 4 | 10 sps | 209,425 | 171,501 | 115,639 | 61.99 | 48.67 | 5.12 |
| 5 | 10 sps | 300,533 | 247,413 | 143,985 | 67.89 | 50.74 | 19.44 |
| 6 | 10 sps | 365,331 | 301,504 | 165,707 | 69.96 | 66.47 | 15.05 |
| 7 | 10 sps | 251,022 | 207,394 | 127,199 | 66.66 | 56.36 | 19.65 |
| 8 | 10 sps | 281,454 | 236,022 | 134,735 | 69.36 | 39.65 | 4.80 |

Table S5: Assessment of COI capture enrichment specificity on two types of DNA templates. "% of COI reads": percentage of reads that align to the COI protein reference database, "% of COI reads assigned to a MC species": percentage of the number of reads that were successfully assigned to a species using the COI nucleotide reference databases to the COI assigned reads and "% of non-protostomian reads": percentage of reads that align to non-protostomian groups. Indicated values are mean number and standard deviation. For sample values, see Table S4 (PCR enrichment) and Table S5 (Capture enrichment). 10 sps: 10 species MC; 52 taxa:52 taxa MC.

**Bulk DNA**

| **Samples** | MC type | Raw | Quality filtered | % reads assigned to COI | % of COI reads assigned to a MC species | % of reads assigned to non-protostomian |
|---|---|---|---|---|---|---|
| 1 | 10 sps | 456,417 | 289,201 | 6.66 | 61.73 | 14.52 |
| 2 | 10 sps | 165,089 | 75,345 | 27.26 | 63.25 | 8.80 |
| 3 | 10 sps | 857,040 | 339,778 | 45.44 | 61.55 | 8.94 |
| 4 | 10 sps | 383,568 | 236,817 | 6.52 | 63.01 | 15.67 |
| 5 | 10 sps | 459,650 | 234,785 | 17.94 | 62.92 | 10.56 |
| 6 | 10 sps | 820,343 | 326,158 | 52.75 | 55.36 | 7.21 |
| 7 | 10 sps | 381,006 | 173,037 | 38.78 | 60.60 | 6.52 |
| 8 | 10 sps | 428,625 | 272,747 | 5.89 | 60.94 | 14.83 |
| A | 52 taxa | 404,000 | 206,457 | 71.99 | 62.46 | 8.71 |
| B | 52 taxa | 424,222 | 201,446 | 73.55 | 64.99 | 5.53 |
| C | 52 taxa | 521,296 | 264,357 | 56.95 | 60.05 | 4.05 |
| D | 52 taxa | 365,939 | 163,938 | 71.24 | 61.09 | 3.75 |
| E | 52 taxa | 501,838 | 234,229 | 69.19 | 62.59 | 6.09 |
| F | 52 taxa | 437,073 | 209,903 | 70.28 | 63.79 | 5.74 |
| G | 52 taxa | 787,464 | 378,592 | 66.10 | 66.62 | 4.29 |
| H | 52 taxa | 643,607 | 298,314 | 75.35 | 64.29 | 6.34 |
| i | 52 taxa | 597,073 | 303,811 | 60.48 | 64.56 | 5.44 |
| J | 52 taxa | 855,743 | 385,902 | 77.90 | 62.46 | 2.69 |

**etDNA**

| **Samples** | MC type | Raw | Quality filtered | % reads assigned to COI | % of COI reads assigned to a MC species | % of reads assigned to non-protostomian |
|---|---|---|---|---|---|---|
| 1 | 10 sps | 575584 | 348292 | 21.93 | 46.65 | 49.14 |
| 2 | 10 sps | 268611 | 150820 | 16.31 | 17.08 | 60.72 |
| 3 | 10 sps | 217240 | 130783 | 12.50 | 35.76 | 69.49 |
| 4 | 10 sps | 451088 | 257570 | 15.21 | 36.38 | 61.68 |
| 5 | 10 sps | 772633 | 506973 | 18.51 | 22.57 | 60.24 |
| 6 | 10 sps | 440802 | 262404 | 13.78 | 31.04 | 59.43 |
| 7 | 10 sps | 500176 | 289496 | 28.05 | 30.81 | 35.94 |
| 8 | 10 sps | 911484 | 530759 | 26.42 | 49.39 | 49.81 |

Table S6: Number of reads recovered for each species in each sample for PCR and capture enrichment and for bulk and ethanol DNA on the 10 species mock communities. Reads were assigned with the blastn algorithm (Camacho et al, 2009). Only alignments with an e-value under $1.10^{-10}$, a query cover over 200 bases for bulk DNA and over 90 bases for etDNA and an identity over 97% were conserved.

| **PCR enrichment** | | | | | | | | | | | |
|---|---|---|---|---|---|---|---|---|---|---|---|
| | Samples | *Chironomus riparius* | *Epeorus assimilis* | *Heptagenia sulphurea* | *Isoperla rivulorum* | *Nemurella picteti* | *Hydropsyche siltalai* | *Athripsodes aterrimus* | *Physella acuta* | *Ancylus fluviatilis* | *Gammarus fossarum* |
| | 1 | 411 | 3838 | 2837 | 1122 | 411 | 2058 | 729 | 231 | 210 | 41 |
| | 2 | 408 | 2074 | 4300 | 1222 | 473 | 2756 | 1035 | 165 | 551 | 6 |
| **Bulk DNA** | 3 | 3674 | 16319 | 24326 | 9227 | 2043 | 24608 | 8501 | 1532 | 4668 | 141 |
| | 4 | 308 | 2595 | 1793 | 710 | 362 | 2283 | 903 | 175 | 603 | 3 |
| | 5 | 1393 | 2538 | 7179 | 2315 | 70 | 8379 | 2788 | 236 | 1597 | 0 |
| **BF2/BR2** | 6 | 3560 | 21675 | 26072 | 7480 | 340 | 22341 | 7396 | 1829 | 4560 | 0 |
| | 7 | 1059 | 12690 | 11915 | 3179 | 144 | 7900 | 1670 | 446 | 1665 | 0 |
| | 8 | 516 | 1593 | 2966 | 920 | 49 | 2685 | 613 | 74 | 380 | 1 |
| | 1 | 335 | 8587 | 30182 | 121 | 6464 | 12287 | 2 | 436 | 1533 | 979 |
| | 2 | 370 | 4416 | 24030 | 2 | 1233 | 9153 | 0 | 355 | 2124 | 807 |
| **etDNA1** | 3 | 163 | 5584 | 18747 | 51 | 9760 | 15954 | 0 | 390 | 1177 | 664 |
| | 4 | 24 | 9054 | 5919 | 0 | 5555 | 30708 | 0 | 43 | 136 | 299 |
| | 5 | 290 | 9056 | 32034 | 7 | 961 | 34095 | 0 | 435 | 2163 | 6185 |
| **fwh1** | 6 | 289 | 766 | 21367 | 0 | 37 | 116177 | 0 | 254 | 1305 | 2 |
| | 7 | 328 | 1640 | 41916 | 330 | 3467 | 29846 | 0 | 96 | 225 | 59 |
| | 8 | 104 | 30227 | 8591 | 33 | 4687 | 20229 | 0 | 32 | 757 | 253 |
| | 2 | 410 | 17950 | 16782 | 0 | 790 | 6121 | 0 | 88 | 382 | 107 |
| | 3 | 343 | 5365 | 15949 | 13 | 1463 | 2718 | 0 | 133 | 921 | 368 |
| **etDNA2** | 5 | 88 | 1280 | 56779 | 0 | 15 | 9145 | 0 | 18 | 176 | 87 |
| | 6 | 724 | 4124 | 5790 | 4 | 128 | 113707 | 0 | 172 | 610 | 52 |
| | 7 | 307 | 4970 | 8221 | 7 | 980 | 61324 | 0 | 121 | 233 | 4 |
| **fwh1** | 8 | 85 | 24863 | 20503 | 25 | 22 | 16647 | 0 | 14 | 197 | 7 |
| | 9 | 242 | 10507 | 15707 | 126 | 20005 | 581 | 2 | 39 | 72 | 8 |
| | 10 | 208 | 18051 | 24550 | 7 | 509 | 9348 | 0 | 89 | 185 | 37 |
| | 2 | 814 | 35327 | 1022 | 55 | 97 | 16 | 190 | 0 | 0 | 3 |
| | 3 | 1648 | 27180 | 2276 | 614 | 857 | 21 | 576 | 0 | 0 | 0 |
| | 5 | 745 | 3228 | 19523 | 49 | 19 | 28 | 62 | 0 | 0 | 0 |
| **etDNA2** | 6 | 6489 | 12788 | 1454 | 377 | 54 | 2815 | 750 | 0 | 0 | 0 |
| | 7 | 1425 | 17016 | 901 | 128 | 9 | 650 | 104 | 2 | 0 | 0 |
| **BF2/BR2** | 8 | 109 | 90009 | 2083 | 51 | 5 | 47 | 6 | 0 | 0 | 0 |
| | 9 | 335 | 5883 | 1841 | 1352 | 6056 | 2 | 5818 | 0 | 0 | 0 |
| | 10 | 373 | 37236 | 11167 | 56 | 120 | 42 | 71 | 0 | 0 | 0 |

| | Capture enrichment | | | | | | | | | | |
|---|---|---|---|---|---|---|---|---|---|---|---|
| | Samples | *Chironomus riparius* | *Epeorus assimilis* | *Heptagenia sulphurea* | *Isoperla rivulorum* | *Nemurella picteti* | *Hydropsyche siltalai* | *Athripsodes aterrimus* | *Physella acuta* | *Ancylus fluviatilis* | *Gammarus fossarum* |
| **Bulk DNA** | 1 | 411 | 3838 | 2837 | 1122 | 411 | 2058 | 729 | 231 | 210 | 41 |
| | 2 | 408 | 2074 | 4300 | 1222 | 473 | 2756 | 1035 | 165 | 551 | 6 |
| | 3 | 3674 | 16319 | 24326 | 9227 | 2043 | 24608 | 8501 | 1532 | 4668 | 141 |
| | 4 | 308 | 2595 | 1793 | 710 | 362 | 2283 | 903 | 175 | 603 | 3 |
| | 5 | 1393 | 2538 | 7179 | 2315 | 70 | 8379 | 2788 | 236 | 1597 | 0 |
| | 6 | 3560 | 21675 | 26072 | 7480 | 340 | 22341 | 7396 | 1829 | 4560 | 0 |
| | 7 | 1059 | 12690 | 11915 | 3179 | 144 | 7900 | 1670 | 446 | 1665 | 0 |
| | 8 | 516 | 1593 | 2966 | 920 | 49 | 2685 | 613 | 74 | 380 | 1 |
| **etDNA** | 1 | 168 | 1552 | 5488 | 1390 | 557 | 412 | 24330 | 539 | 1185 | 4 |
| | 2 | 166 | 363 | 1348 | 13 | 83 | 244 | 269 | 566 | 1147 | 2 |
| | 3 | 121 | 144 | 1759 | 152 | 1026 | 481 | 384 | 678 | 1078 | 26 |
| | 4 | 203 | 2926 | 3545 | 28 | 2918 | 3377 | 318 | 405 | 524 | 8 |
| | 5 | 640 | 1743 | 6211 | 224 | 38 | 3919 | 1012 | 2200 | 5185 | 14 |
| | 6 | 268 | 109 | 2327 | 6 | 18 | 6505 | 155 | 359 | 1477 | 0 |
| | 7 | 998 | 505 | 14337 | 2845 | 791 | 3491 | 739 | 363 | 956 | 0 |
| | 8 | 858 | 37178 | 8248 | 565 | 4540 | 7602 | 373 | 538 | 9359 | 5 |
| **Enrichment-free** | 2 | 0 | 0 | 0 | 1 | 1 | 1 | 4 | 0 | 0 | 0 |
| | 4 | 0 | 1 | 1 | 1 | 2 | 2 | 3 | 0 | 0 | 0 |
| | 5 | 1 | 19 | 13 | 2 | 6 | 49 | 5 | 3 | 21 | 0 |
| | 8 | 0 | 1 | 5 | 0 | 2 | 7 | 4 | 1 | 0 | 0 |

Table S7: Number of reads recovered for each taxon in each sample for PCR and capture enrichment for bulk DNA on the 52 taxa mock communities. Reads were assigned with the blastn algorithm (Camacho et al, 2009). Only alignments with an e-value under 1.10-10, a query cover over 200 bases for bulk DNA and an identity over 97% were conserved.

| PCR enrichment | | | | | | | | | | |
|---|---|---|---|---|---|---|---|---|---|---|
| **Taxa** | **A** | **B** | **C** | **D** | **E** | **F** | **G** | **H** | **i** | **J** |
| *Agabus* | 8 | 34 | 109 | 4 | 0 | 33 | 2 | 4 | 5 | 95 |
| *Agapetinae* | 123 | 375 | 0 | 87 | 20 | 323 | 133 | 215 | 25 | 41 |
| *Ancylus* | 0 | 0 | 0 | 0 | 0 | 2 | 0 | 0 | 0 | 0 |
| *Anomalopterygella chauvinia* | 392 | 253 | 66 | 71 | 272 | 558 | 190 | 201 | 215 | 155 |
| *Arhynchobdellida* | 345 | 227 | 260 | 50 | 210 | 258 | 90 | 10 | 352 | 9 |
| *Asellus aquaticus* | 306 | 178 | 140 | 49 | 5681 | 2430 | 618 | 263 | 915 | 168 |
| *Baetis* | 554 | 3175 | 108 | 188 | 218 | 3474 | 526 | 494 | 1261 | 349 |
| *Bivalvia* | 12 | 4 | 0 | 0 | 2 | 10 | 0 | 4 | 0 | 0 |
| *Blephariceridae* | 1671 | 3670 | 310 | 250 | 1402 | 1348 | 543 | 366 | 782 | 154 |
| *Ceratopogonidae* | 284 | 210 | 176 | 209 | 165 | 1036 | 55 | 310 | 60 | 100 |
| *Chaetopteryx villosa* | 334 | 245 | 127 | 56 | 68 | 27 | 76 | 48 | 30 | 130 |
| *Chironomidae* | 763 | 135 | 172 | 32 | 1032 | 1402 | 304 | 380 | 0 | 8 |
| *Cordulegaster boltonii* | 29 | 13 | 32 | 27 | 81 | 295 | 81 | 89 | 62 | 44 |
| *Daphnia pulex* | 9 | 7 | 0 | 0 | 3 | 11 | 7 | 0 | 3 | 4 |
| *Dicranota* | 66 | 9 | 22 | 0 | 3 | 63 | 13 | 25 | 9 | 6 |
| *Drusus annulatus* | 340 | 227 | 112 | 32 | 216 | 611 | 209 | 156 | 129 | 105 |
| *Dugesia* | 25 | 45 | 28 | 14 | 40 | 160 | 30 | 51 | 45 | 19 |
| *Ecdyonurus* | 222 | 438 | 4848 | 101 | 485 | 5346 | 2448 | 96 | 214 | 14696 |
| *Epeorus* | 1444 | 788 | 24613 | 361 | 932 | 1813 | 1892 | 731 | 2782 | 4384 |
| *Ephemerella (Torleya) major* | 207 | 1970 | 1214 | 1453 | 2169 | 843 | 8644 | 7155 | 4177 | 1173 |
| *Ephemerella mucronata* | 16 | 137 | 68 | 15 | 68 | 451 | 105 | 194 | 60 | 30 |
| *Ephemeridae* | 86 | 3407 | 2364 | 798 | 221 | 2466 | 106 | 134 | 1137 | 2054 |
| *Gammarus* | 50 | 15 | 40 | 8 | 46 | 353 | 110 | 54 | 145 | 24 |
| *Gerroidea* | 6617 | 699 | 444 | 513 | 707 | 2746 | 4354 | 3336 | 1723 | 361 |
| *Halesus* | 135 | 64 | 198 | 190 | 87 | 411 | 136 | 182 | 335 | 223 |
| *Hydropsyche* | 6 | 28 | 15 | 5 | 38 | 177 | 41 | 42 | 31 | 26 |
| *Isoperla* | 48 | 332 | 225 | 184 | 522 | 1038 | 592 | 554 | 582 | 254 |
| *Leptophlebiidae* | 544 | 1225 | 274 | 150 | 513 | 3105 | 693 | 3602 | 4281 | 1006 |
| *Leuctra* | 531 | 691 | 534 | 988 | 521 | 5155 | 471 | 513 | 126 | 315 |
| *Limnius* | 174 | 7 | 19 | 17 | 21 | 308 | 50 | 23 | 2 | 5 |
| *Limoniidae* | 166 | 109 | 20 | 19 | 29 | 166 | 75 | 50 | 99 | 13 |
| *Lymnaeidae* | 7 | 0 | 0 | 0 | 0 | 6 | 0 | 0 | 2 | 0 |
| *Nemouridae* | 3909 | 941 | 41 | 240 | 2933 | 4677 | 4818 | 3027 | 647 | 246 |
| *Odontocerum albicorne* | 543 | 302 | 252 | 23 | 170 | 677 | 348 | 164 | 231 | 135 |
| *Oecismus monedula* | 2882 | 454 | 262 | 111 | 375 | 1700 | 856 | 268 | 336 | 258 |
| *Oreodytes sanmarkii* | 2354 | 727 | 2363 | 681 | 2696 | 4021 | 9409 | 6883 | 15257 | 5078 |
| *Perlidae* | 1077 | 478 | 1219 | 1342 | 2022 | 1851 | 616 | 1238 | 1007 | 605 |

| | | | | | | | | | | |
|---|---|---|---|---|---|---|---|---|---|---|
| *Philopotamidae* | 49 | 16 | 9 | 6 | 30 | 124 | 19 | 46 | 24 | 34 |
| *Polycentropodidae* | 0 | 13 | 12 | 0 | 0 | 48 | 79 | 30 | 23 | 17 |
| *Potamophylax* | 186 | 1386 | 223 | 109 | 282 | 1472 | 694 | 470 | 150 | 108 |
| *Ptychoptera* | 119 | 101 | 146 | 54 | 83 | 526 | 196 | 107 | 58 | 70 |
| *Rhithrogena* | 136 | 641 | 1224 | 427 | 1347 | 3184 | 729 | 1506 | 1214 | 6 |
| *Rhyacophila* | 97 | 71 | 44 | 16 | 76 | 374 | 31 | 15 | 62 | 15 |
| *Scirtidae* | 8718 | 2882 | 2055 | 2212 | 2244 | 3879 | 955 | 1877 | 5143 | 260 |
| *Sericostoma personatum* | 695 | 1163 | 509 | 428 | 297 | 1314 | 1028 | 598 | 718 | 542 |
| *Sialis* | 44 | 41 | 71 | 184 | 295 | 1419 | 185 | 158 | 868 | 707 |
| *Silo* | 57 | 25 | 72 | 41 | 279 | 547 | 96 | 2 | 473 | 111 |
| *Simuliidae* | 80 | 68 | 29 | 2 | 15 | 177 | 4 | 72 | 23 | 23 |
| *Thremma gallicum* | 144 | 136 | 132 | 152 | 253 | 65 | 125 | 15 | 54 | 11 |
| *Tipulidae* | 271 | 166 | 114 | 121 | 61 | 901 | 248 | 41 | 175 | 56 |
| *Trombidiformes (Acari)* | 104 | 73 | 71 | 56 | 114 | 327 | 93 | 93 | 70 | 5 |

**Capture enrichment**

| **Taxa** | **A** | **B** | **C** | **D** | **E** | **F** | **G** | **H** | **i** | **J** |
|---|---|---|---|---|---|---|---|---|---|---|
| *Agabus* | 151 | 3341 | 5449 | 125 | 433 | 368 | 249 | 717 | 511 | 8472 |
| *Agapetinae* | 2809 | 1926 | 0 | 1375 | 763 | 1585 | 1699 | 2617 | 1532 | 1311 |
| *Ancylus* | 1840 | 814 | 585 | 174 | 856 | 620 | 799 | 951 | 245 | 445 |
| *Anomalopterygella chauvinia* | 671 | 758 | 565 | 506 | 1131 | 521 | 896 | 1179 | 1025 | 1330 |
| *Arhynchobdellida* | 826 | 619 | 1561 | 243 | 1451 | 190 | 1268 | 115 | 2185 | 242 |
| *Asellus aquaticus* | 7267 | 4036 | 4456 | 3549 | 3995 | 5734 | 13119 | 8695 | 12719 | 10836 |
| *Baetis* | 1937 | 9326 | 1301 | 1914 | 1439 | 3944 | 3336 | 4001 | 4143 | 6070 |
| *Bivalvia* | 531 | 242 | 179 | 51 | 86 | 109 | 23 | 1013 | 167 | 50 |
| *Blephariceridae* | 3487 | 4579 | 1926 | 1710 | 3497 | 1585 | 2367 | 2478 | 2620 | 2106 |
| *Ceratopogonidae* | 355 | 89 | 154 | 451 | 303 | 308 | 1427 | 486 | 250 | 713 |
| *Chaetopteryx villosa* | 1056 | 1137 | 1338 | 611 | 495 | 82 | 880 | 787 | 442 | 2142 |
| *Chironomidae* | 3694 | 1526 | 2201 | 320 | 4547 | 3841 | 3746 | 5078 | 528 | 4796 |
| *Cordulegaster boltonii* | 735 | 273 | 1158 | 1140 | 2620 | 2484 | 3403 | 4104 | 2598 | 3120 |
| *Daphnia pulex* | 5 | 1 | 4 | 1 | 1 | 1 | 0 | 0 | 0 | 0 |
| *Dicranota* | 815 | 223 | 728 | 215 | 185 | 520 | 1066 | 757 | 406 | 486 |
| *Drusus annulatus* | 738 | 861 | 944 | 328 | 1311 | 776 | 1873 | 1735 | 1181 | 2424 |
| *Dugesia* | 0 | 0 | 0 | 0 | 0 | 0 | 4 | 0 | 0 | 0 |
| *Ecdyonurus* | 1361 | 3236 | 601 | 1445 | 4083 | 345 | 432 | 1933 | 1954 | 2274 |
| *Epeorus* | 574 | 290 | 5879 | 265 | 510 | 455 | 521 | 571 | 732 | 5046 |
| *Ephemerella (Torleya) major* | 219 | 1958 | 2092 | 1725 | 2007 | 353 | 4885 | 5415 | 2720 | 3739 |
| *Ephemerella mucronata* | 236 | 2703 | 1846 | 734 | 2254 | 1593 | 2409 | 5252 | 1356 | 2599 |
| *Ephemeridae* | 274 | 3859 | 4127 | 2556 | 800 | 2818 | 848 | 1019 | 2532 | 11004 |
| *Gammarus* | 328 | 227 | 624 | 132 | 632 | 621 | 1075 | 1091 | 1357 | 937 |
| *Gerroidea* | 7833 | 1859 | 2109 | 2159 | 2597 | 3472 | 11773 | 11183 | 4865 | 3251 |
| *Halesus* | 748 | 991 | 1831 | 854 | 914 | 726 | 1236 | 1781 | 1662 | 2807 |
| *Hydropsyche* | 3170 | 1247 | 3008 | 1698 | 734 | 937 | 5741 | 4677 | 608 | 6350 |
| *Isoperla* | 371 | 2113 | 1874 | 1565 | 3084 | 1463 | 4778 | 6121 | 2852 | 4591 |

| | | | | | | | | | | |
|---|---|---|---|---|---|---|---|---|---|---|
| *Leptophlebiidae* | 1459 | 4139 | 1522 | 1713 | 2264 | 1781 | 3848 | 4616 | 10116 | 10286 |
| *Leuctra* | 1398 | 2119 | 2990 | 3742 | 2087 | 6721 | 2728 | 5210 | 1040 | 4361 |
| *Limnius* | 1576 | 277 | 414 | 747 | 262 | 888 | 1474 | 899 | 273 | 823 |
| *Limoniidae* | 1422 | 1191 | 506 | 704 | 877 | 731 | 1223 | 846 | 793 | 775 |
| *Lymnaeidae* | 2112 | 1544 | 405 | 850 | 1155 | 1672 | 984 | 1476 | 1933 | 415 |
| *Nemouridae* | 9675 | 5020 | 1655 | 5883 | 10129 | 6112 | 6591 | 16068 | 3186 | 4872 |
| *Odontocerum albicorne* | 5699 | 5470 | 7248 | 2492 | 5915 | 4751 | 9563 | 4745 | 4733 | 8410 |
| *Oecismus monedula* | 2314 | 1286 | 1347 | 688 | 1357 | 1453 | 2880 | 1704 | 1646 | 2800 |
| *Oreodytes sanmarkii* | 3101 | 2153 | 2108 | 1000 | 978 | 4052 | 10687 | 4615 | 7555 | 9667 |
| *Perlidae* | 614 | 348 | 456 | 2709 | 2257 | 627 | 636 | 861 | 437 | 734 |
| *Philopotamidae* | 6080 | 2416 | 3397 | 5190 | 5376 | 4442 | 4982 | 6736 | 3949 | 13881 |
| *Polycentropodidae* | 143 | 2366 | 3220 | 707 | 93 | 2404 | 10300 | 6222 | 2970 | 4956 |
| *Potamophylax* | 209 | 1707 | 667 | 280 | 722 | 1139 | 1669 | 1672 | 1021 | 1132 |
| *Ptychoptera* | 1314 | 2139 | 6760 | 3532 | 3631 | 2781 | 8030 | 4719 | 2230 | 7487 |
| *Rhithrogena* | 388 | 1702 | 3360 | 1794 | 2877 | 1676 | 3265 | 2289 | 2425 | 5984 |
| *Rhyacophila* | 1580 | 578 | 1055 | 730 | 1139 | 1101 | 841 | 733 | 806 | 1617 |
| *Scirtidae* | 3237 | 2971 | 3747 | 2398 | 1740 | 1441 | 3282 | 2652 | 3215 | 3315 |
| *Sericostoma personatum* | 363 | 651 | 579 | 473 | 253 | 336 | 944 | 761 | 582 | 1363 |
| *Sialis* | 339 | 1301 | 2331 | 3421 | 4012 | 4720 | 3742 | 4589 | 5986 | 15964 |
| *Silo* | 2120 | 2137 | 2448 | 2387 | 8151 | 5953 | 3442 | 668 | 9485 | 4623 |
| *Simuliidae* | 2339 | 1335 | 937 | 689 | 985 | 800 | 2048 | 2244 | 1063 | 1362 |
| *Thremma gallicum* | 2384 | 964 | 3499 | 1599 | 1677 | 527 | 5659 | 1107 | 848 | 1272 |
| *Tipulidae* | 464 | 360 | 460 | 461 | 150 | 607 | 702 | 197 | 482 | 456 |
| *Trombidiformes (Acari)* | 143 | 131 | 178 | 100 | 183 | 163 | 272 | 363 | 168 | 380 |

Figure S1: Robustness of PCR (top) and capture (bottom) enrichment to mismatches between the targeted species COI and the best pair of degenerated primers or the best bait.

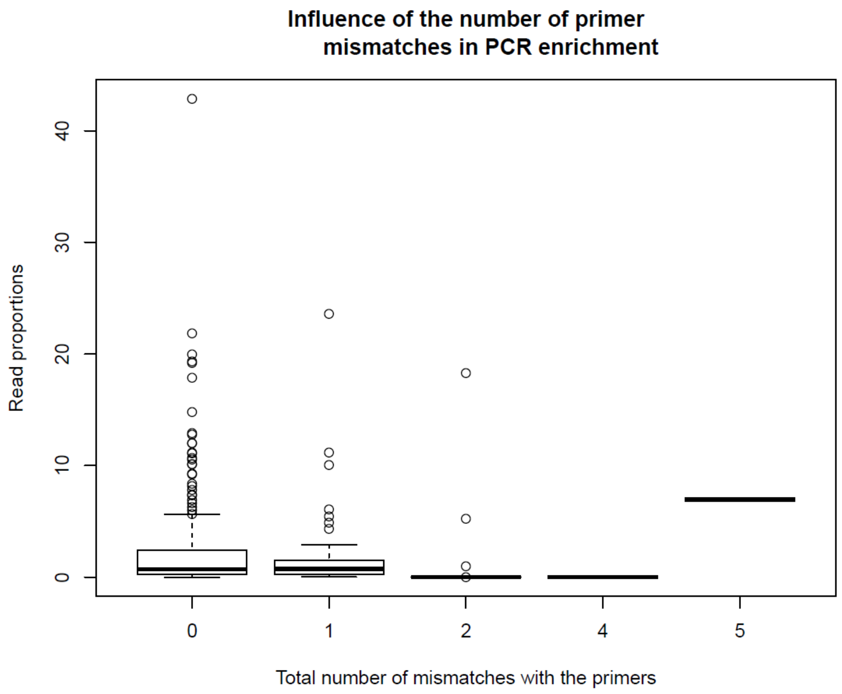


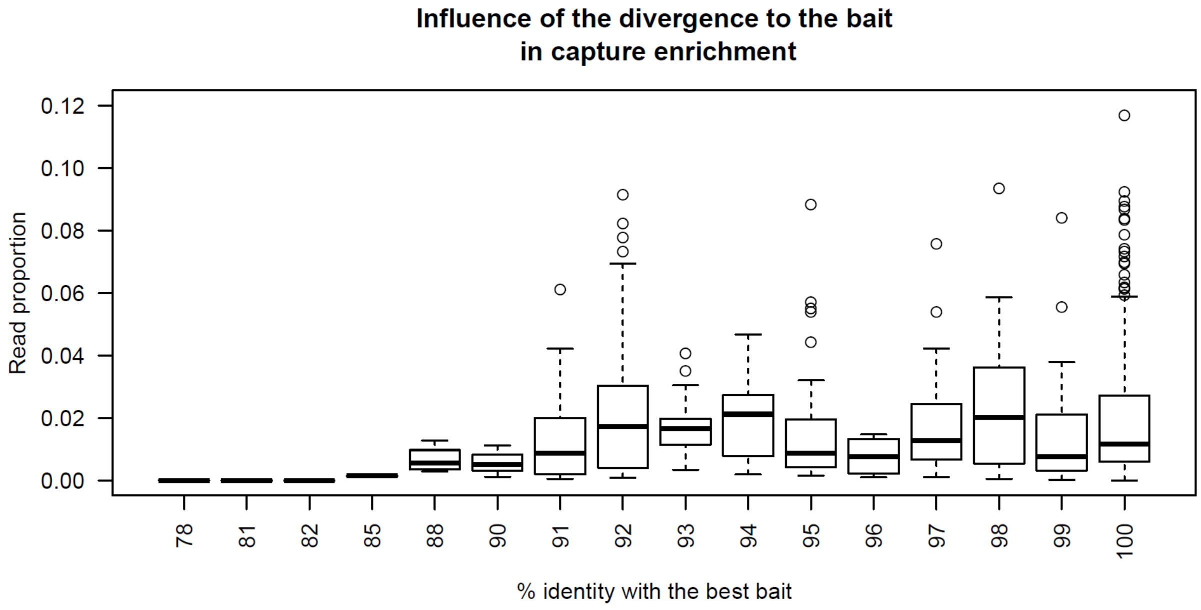

Figure S2: Percentage of reads for different coarse taxonomic group (archaea, eubacteria, fungi, plant, protist, protostomia (here Arthropoda, Annelida, Brachiopoda, Mollusca, Nematoda and Platyhelminthes) and other metazoa (corresponding to deuterostomia)) for each mock community type (10 sps = 10 species MC; 52 taxa=52 taxa MC), template DNA (bulk DNA and etDNA) and enrichment method (capture enrichment and PCR enrichment (BF2/BR2 primer pair for bulk DNA and fwh1 primer pair for etDNA)). The assignment was obtained by comparing the quality filtered reads to a protein database designed using diamond (blastx, more sensitive option, e-value threshold of $1.10^{-10}$) (Buchfink et al, 2015).

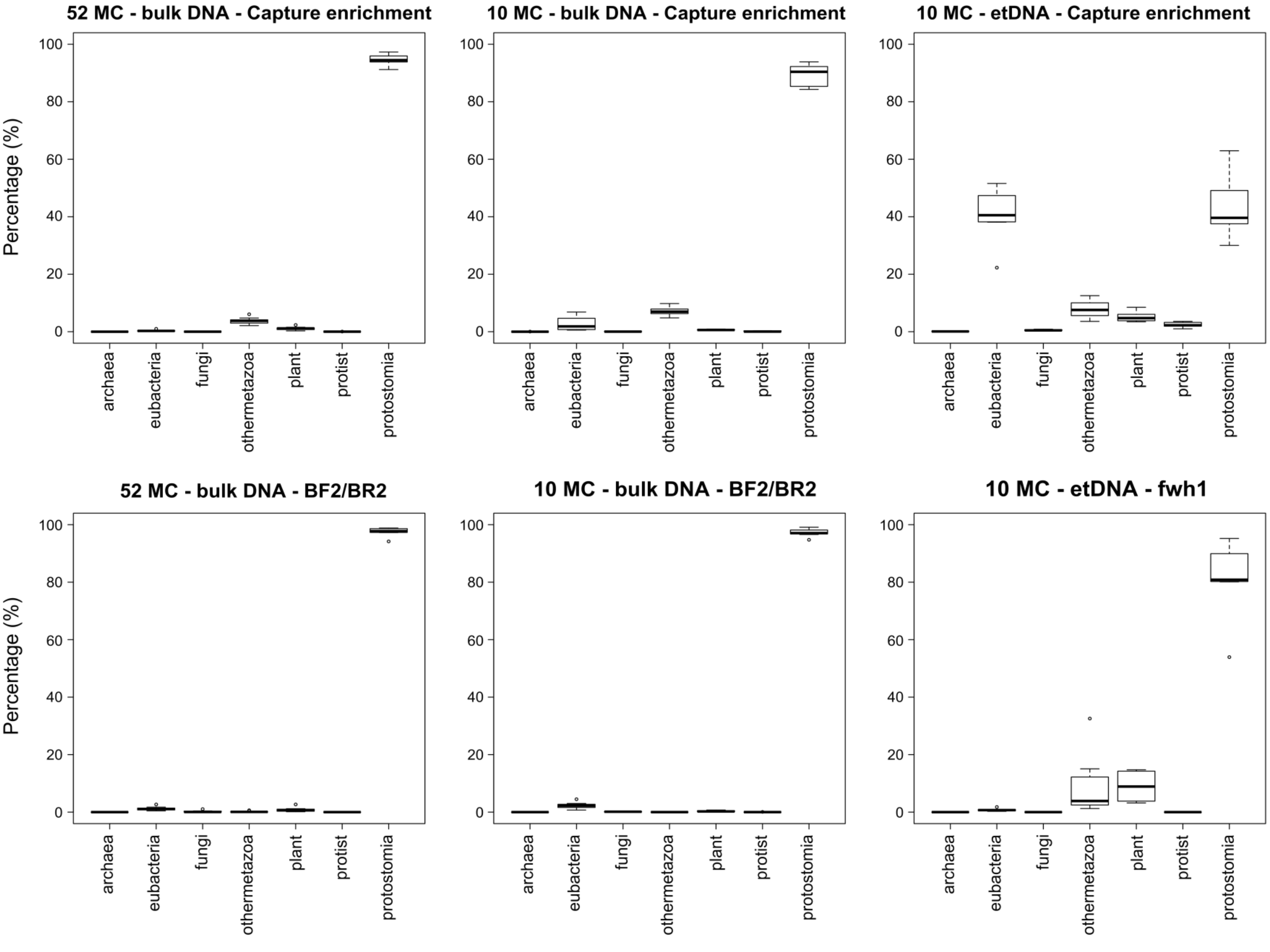
